# A quantitative spatial atlas of transcriptomic, morphological, and electrophysiological cell type densities in the mouse brain

**DOI:** 10.1101/2024.12.30.630728

**Authors:** Csaba Verasztó, Yann Roussel, Lida Kanari, Henry Markram, Daniel Keller

## Abstract

Brain cells can be classified into transcriptomic, morphological, and electrophysiological types. We derived a quantitative spatial atlas of the distributions of these classes within the mouse brain. To do so, we first generated a 3D atlas of transcriptomic cell type densities, by scaling regional estimates of densities from brain slices using cell counts and slice dimensions. Adjustments for regions with high cell density, such as the cerebellum, were made using the average Nissl intensities as a density modifier on a voxel-by-voxel basis. To connect the transcriptomic-type signatures to cellular function (morphological-electrophysiological types), we leveraged patch-sequencing datasets, which integrate mRNA counts, morphology reconstructions, and electrophysiological recordings from single cortical neurons. We aligned mRNA counts with the established classifications of transcriptomic types, classified neuron morphologies according to morphological types, and derived electrophysiological types using K-means clustering. The resulting 3D atlas and computational framework can be used to probe neuronal cell-type diversity and its functional implications, setting the stage for further explorations into the cellular basis of brain function.

## Introduction

While neural networks are well-characterized in a few invertebrate species [1–4], the mouse brain — with its 70 million neurons [5] — can serve as a bridge between simpler organisms and more complex vertebrate nervous systems, such as humans, in neuroscience research. For the mouse brain, current initiatives emphasize whole-brain approaches [6–9] to decipher the vast cellular diversity within it. These and other ambitious efforts [10–12] have yielded insights into the brain’s complex organization and function. Over the past decade, single-cell RNA sequencing (scRNA-seq) [13–15] and single-nucleus RNA sequencing [16–18] revealed how gene expression varies across different brain regions [19–21]. Recently, comprehensive sampling of the entire mouse brain, often with single-molecule precision resolution [22,23] and within a common platform [24], now allows high-resolution mapping of neural cell types across the brain. The integration of large amounts of spatial transcriptomic data [9,25] in the whole-brain broader context, revealed complex interregional relationships, and precise analysis of tissue characteristics. However, since no mouse brain has yet been fully mapped at the single-cell level, estimations of transcriptomic cell type placement will remain a necessary tool for researchers until complete cellular identification can be realized on the whole-brain or whole-body scale.

Using transcriptomic data [8,9] and the common coordinate framework from the Allen Institute for Brain Science (AIBS) [25], we created 3D atlases that map the spatial distribution of distinct transcriptomic cell types (t-types). We extracted cell coordinates and their brain region’s volume from aligned sections to estimate cell densities across all brain regions. As sections often provide multiple samples from a region, coverage of the mouse brain is well-validated and almost complete.

Additionally, we developed a method for converting between various cell type classification systems. Recent advances in patch-sequencing technologies link transcriptomic profiles of individual cells with morphological types (m-types), which categorize cell shape and structure, and electrophysiological types (e-types), which categorize cells according to electrophysiological behavior [26–28]. From such experimental data, we derived a model to convert between various cell type classifications and recreate the 3D atlas for all other functional m- and e-type classifications.

Hitherto, much scientific effort has focused on mapping the densities of inhibitory and excitatory neurons in the brain. The methods developed in this paper go beyond this by integrating cell type modalities (t-, m- and e-) on a whole-brain scale. Converting transcriptomic cell clusters into previous functional morphological-electrophysiological classifications (me-classifications) bridges the gap with classifications present in the traditional body of literature, allowing quantitative comparison of older cell counting results with transcriptomic results. This provides neuroscientists with an accurate cell density map of the mouse brain that they can easily interpret.

## Methods

### Data used for models

In this study, we combined several whole-brain datasets to provide a more robust and detailed perspective: The Allen Brain Cell (ABC) Atlas [24], the multiplexed error-robust fluorescence *in situ* hybridization (MERFISH) dataset [19,24], the extended version of the latest common coordinate framework annotation (CCFv3) and Nissl volume [29] from the AIBS and the Blue Brain Project, and two patch-seq datasets [27,28] (Figure 1A). The ABC Atlas is an online platform that provides a comprehensive cell type atlas, visualizing transcriptomic cell type expression, spatial organization, and diversity across the brain. MERFISH is a high-resolution spatial transcriptomics technique that quantifies and maps RNA molecules within cells. For our analysis, we downloaded both the complete mouse brain taxonomy (hierarchical clustering) and MERFISH dataset from the platform. All analyses were conducted using the 2024-03-30 release of the ABC Atlas to ensure consistency and alignment with the latest available data. Patch-seq datasets are resourceful because they link RNA sequencing, morphological reconstruction and electrophysiological recordings for the same neuron type. Each used dataset focuses on a different region of the mouse cortex: primary visual cortex for Gouwens et al. [27] (∼500 inhibitory neurons) and primary motor cortex for [28] Scala et al. (∼200 pyramidal cells and ∼300 inhibitory neurons).

**Figure 1:**
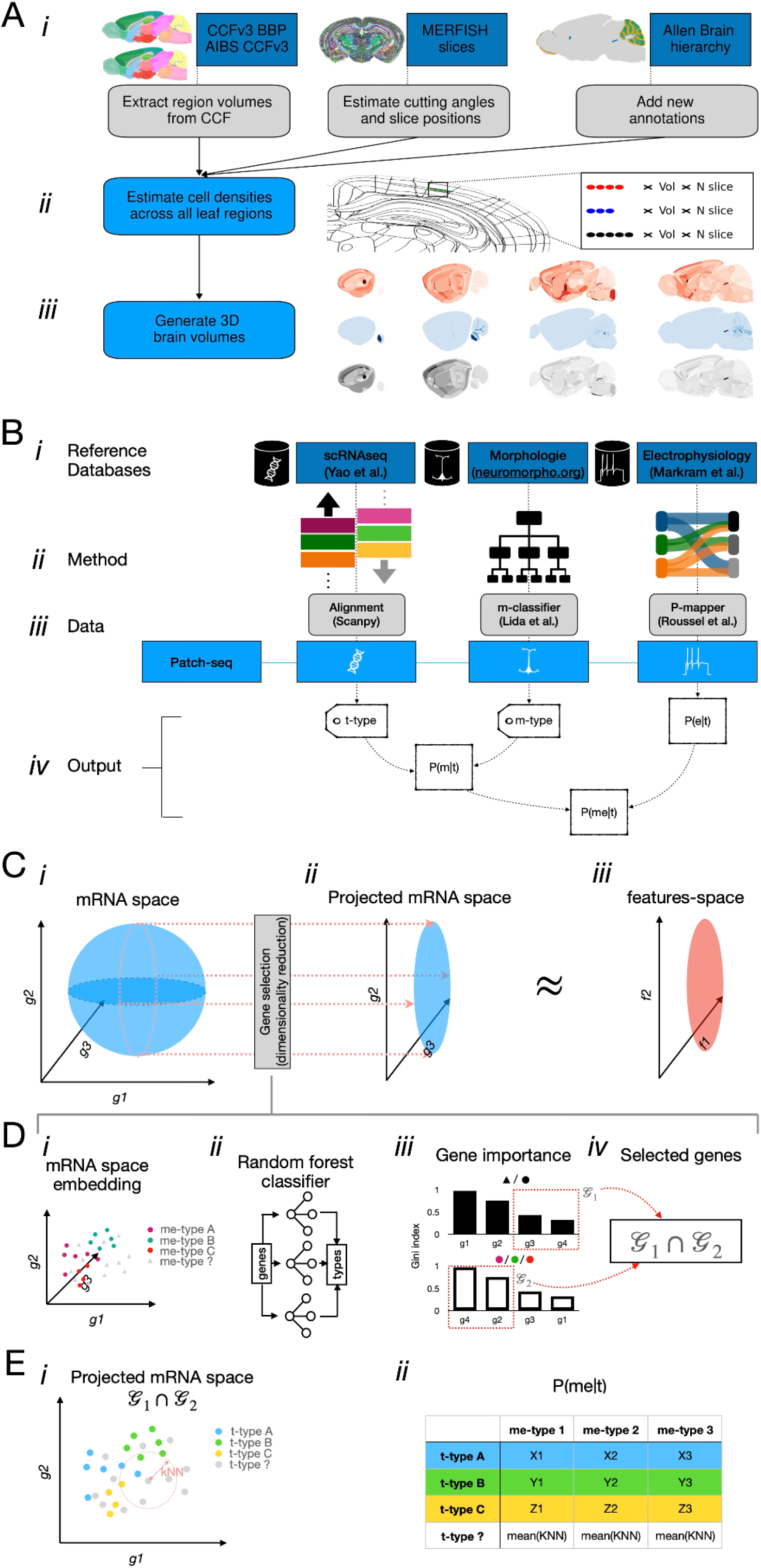
Pipeline overview of probabilistic map P(me|t) computation. **A.** Pipeline to generate 3D transcriptomic cell type atlases from MERFISH slices. We produced both unscaled and scaled versions of the densities by (i) integrating various datasets with MERFISH metadata, (ii) calculating cell-type densities for each leaf region of the mouse brain, and (iii) projecting these densities back into the average brain space. **B.** The patch-seq data derived P(me|t) is derived in four steps. Reference dataset for the three modalities (*i*) were used to align (*ii*) the patch-seq dataset (*iii*) on established classification and derive the probability of observing an me-type given a t-type (*iv*) **C.** Extension of P(me|t) rely on a dimensionality reduction of the mRNA space so that the resulting projected space mirrors at best the feature space. **D.** Detailed overview of the gene selection process for the projected mRNA space. **E.** Probabilities of uncovered t-types are approximated using the averaged probabilities of the neighboring covered t-types on the projected mRNA space using kNN.

### Extending CCFv3’s hierarchical annotations and parcellations for comprehensive dataset integration

We extended the hierarchical annotation of the AIBS CCFv3 annotation volume [25] for the whole dataset in which all voxels could be assigned to a leaf region (the term leaf region represents the deepest level of the mouse brain ontology). We also extended the parcellation to parcellation term membership metadata that was created for the ABC Atlas. These steps were necessary to resolve inconsistencies between the original structural annotations of the CCF and the ABC Atlas parcellation system. We incorporated label IDs and alternative cluster names, omitting special symbols to facilitate easier processing. Finally, the *main olfactory bulb, glomerular laye*r was added to the list as a substructure, as well as all granular, molecular, and Purkinje layers of all the lobules of the cerebellum. In all steps, we adhered to the ABC Atlas’s new multimodal reference system which introduced parcellations, while maintaining backward compatibility with the CCFv3’s hierarchical annotations.

### Aligning and resampling brain annotation volumes for improved cell density estimation

The ABC Atlas provides an aligned and resampled annotation volume in which region IDs were renumbered for better visualization. However, it is not perfectly aligned with the CCFv3 annotation volume. Therefore, we used the original CCFv3 annotation volume (at 10 μm voxel resolution) and identified all original region IDs by matching voxel counts in the corresponding annotation volume slice for every cell. This gave access to a rotated, resampled and aligned annotation array with exact volumes for every brain region intersected by MERFISH slices (Fig. 1 A*i middle*). Consequently, cells of every cell type located in these brain regions acquired the exact brain volume of their brain region within the brain slices as metadata. Acquiring this information allowed us to overcome issues like estimating cell type densities in multiple patches (e.g. a brain region was intersected in both hemispheres by the same MERFISH slice), or estimating cell densities by triangulating region volumes around their CCF coordinates.

### Estimation of cutting angles and coronal region volumes for MERFISH sections

For the coronal MERFISH sections, cutting angles (tilted in both horizontal and vertical directions) and coronal positions were estimated using non-mutual information (NMI) metrics [30], aided by the provided average template array. For all 53 coronal slices the estimated axial angles were 36.84 ± 3.3 degrees, while the vertical angles were 83.64 ± 0.27 degrees. This information was used to calculate the region’s volume of the MERFISH section in the plane for each individual cell.

For every cell in the ABC Atlas database next to its single reconstructed z coordinate (i.e. along the coronal axis), we added their voxel position, their template number, cutting angles , the region ID of their leaf region, together with the region’s voxel count (i.e. voxelized volume) (Fig. 1 A*i*). Cells which did not pass the ABC Atlas quality control (QC) were not included in the calculations. Additionally, 15 cells were removed from the database because their incorrect x_ccf and y_ccf coordinates were 0. Although these cells passed the ABC Atlas QC, we could not use them since they were physically located outside of the brain. It is important to note that artifacts from imperfect sectioning, tissue damage, or chemical processing of the slices could obscure the estimated volumes. However, many of these irregularities could be resolved or corrected during the registration and alignment of MERFISH sections to the common coordinate framework. We estimated volumes from the annotation volume file (Fig. 1 A*i* left) registered to the MERFISH slices and incorporated this data into the metadata for each cell. Missing tissue, bad quality cell information or cells registered to uncharacterised regions could not be recovered.

After the preceding steps, 3,791,571 cells remained. These cells belonged to 677 regions in a total of 53 coronal sections (Fig. 1 A*ii*). Out of these regions, the newly added main olfactory bulb (MOB) layer and the cerebellar lobe layers remained empty, as they were either not part of the ABC’s parcellation annotations or no cells were identified in the region which passed QC. We split these substructures into their respective leaf regions (Fig. 1 A*i* right) based on a simple set of rules. Cerebellar lobules can be split into Purkinje, molecular, and granular layers. Purkinje layers inherited 100% of the Purkinje cells, 100% of the Bergman glia, and they shared microglia equally with the other two leaf regions. Molecular layers were set to be purely inhibitory, thus they do not contain any Bergmann glia, Purkinje cells or cells expressing the neurotransmitter glutamate (Glut). Granular layers inherited every cell type, except Purkinje cells and Bergmann glia; and only contain Golgi cells from the gamma-aminobutyric acid (GABA) expressing cell types. On the other hand, the missing *main olfactory bulb, glomerular laye*r has inherited all the cells from the unassigned region of the MOB.

It is important to mention that MERFISH slices did not completely cover every brain region: e.g. the *frontal pole, layer 1*, and retrosplenial area, dorsal part, layer 4 were not covered in this dataset. We included Supplementary Table 1 listing all leaf regions with no direct coverage. Additionally, a couple substructures covering white matter of the brain were not broken down to their respective leaf regions in the AIBS’s parcellation annotations; thus they were used instead, which gives these areas a more crude resolution. These regions are: *corpus callosum*, *anterior forceps* with two regions, the inferior *cerebellar peduncle* with two regions, the *sensory root of the trigeminal nerve* with two regions, the *superior cerebellar peduncles* with four regions, and the *stria terminalis* with two regions. These regions inherited the same set of cell types and densities. We also recognized that zero density values can convey valuable biological information; thus, for regions without assigned voxels in the 3D reference space, we used N/A (not a number) to distinguish between the lack of information and the absence of cells.

Cells which were assigned to unassigned brain regions in the ABC Atlas were omitted from the calculations. This omission did not influence regional cell densities but reduced somewhat the overall cell count. Note that we have only included cells from the ABC Atlas where the CCF positional coordinates were identified, as the alignment process of MERFISH sections were not available to us. To reduce data processing and avoid potential bleed over of cells from neighboring regions due to misalignment, we decided to not include sections which were not aligned to the common coordinate framework. Instead we rigorously tested the ABC Atlas alignment ourselves to check for misalignment which could be detected in regions where cell composition was known from the literature.

### Estimation of cell type densities in their respective region

Densities were estimated where alignment to the common coordinate framework was available, with volumes calculated by multiplying the voxelized region size in each slice by the section thickness (10 µm in the fresh-frozen state). Densities were estimated on a cell type by region basis by summing all cells and dividing by the region’s total volume across all intersecting slices. (Fig 2). MERFISH slices cover 565 brain regions (parcellation substructures) where cell types were identified and passed the QC. Following the annotation hierarchy, these regions were divided into their respective leaf regions, resulting in cell coverage for 670 of the 677 possible regions. Out of the 7 missing regions (*frontal pole, layer 1, central canal, spinal cord/medulla, pyramidal decussation, bed nucleus of the anterior commissure, cuneate fascicle, vomeronasal nerve, Gracile nucleus*) only the first is from the gray matter. Brain regions contained on average 4819 (median cell count was only 1430) cells, corresponding to 124 cell types (neuron and non-neuron types together). On average, leaf regions were crossed by a slice 7.13 times (median was 5). By splitting cerebellar regions into their respective leaf regions, these values were slightly different (as 16 lobules are split into 16 times 3 leaf regions). For the seven missing regions, we kept cell type numbers as N/A, but it was possible to approximate densities using scaling. However a more accurate estimate could be achieved by incorporating additional MERFISH slices into the current pipeline.

**Figure 2:**
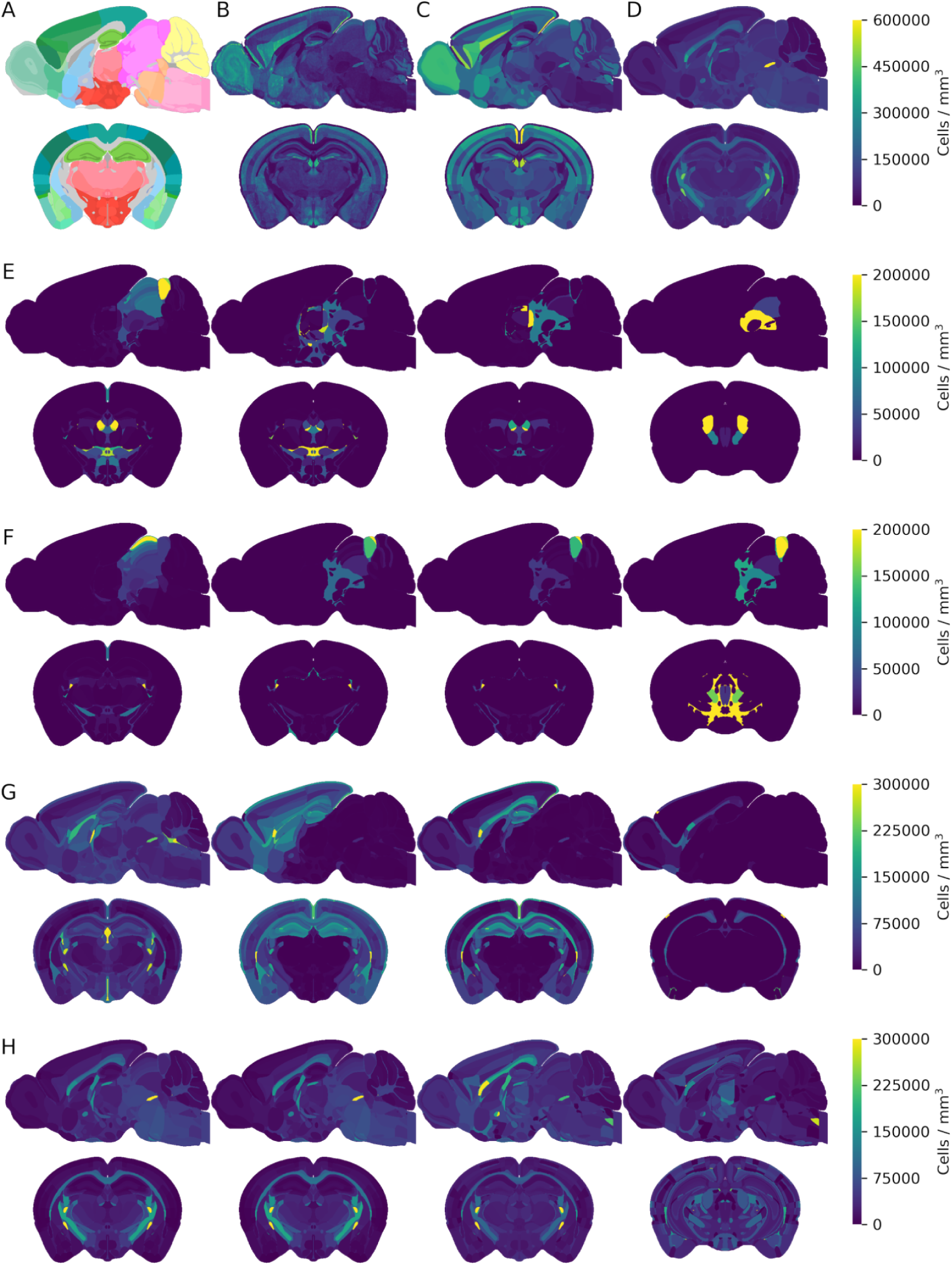
Regional transcriptomic cell counts in 3D atlas form (528, 320, 456). Sagittal (y=200) and coronal (x=300) sections from the **A.** extended CCFv3 annotation volume, **B.** Regional cell counts with Nissl granularity, **C.** Regional neuron counts (averages), and **D.** regional non-neuronal counts (averages). Colorbar shows cell numbers in number of cells / mm^3^ for panel B. **E.** Sagittal (y=200) and coronal (x=300) sections of an excitatory cell type at 4 different hierarchies (class, subclass, supertype, cluster). **F**. Inhibitory cell type examples at different hierarchical levels. **G.** Astrocyte type examples at different hierarchical levels. **H.** Oligodendrocyte cell type examples at different hierarchical levels. Colorbars (**E-H**) show cell numbers for the first cell type in the row.

### Cell density estimation and scaling methods

After densities were estimated across the whole brain, we leveraged several literature values to improve the initial estimations. To achieve this we developed three separate methods to improve estimates: global scaling, cell type density transplant, and Nissl staining based estimation (Fig. 2S2). Users can choose which method to employ based upon their needs. Using voxel-based volumetric information, we can calculate the total cell numbers for every cell type or cell groups and scale up/down the densities in the regions.

In the global scaling method, we first pooled all 5,274 cell types present in MERFISH slices in larger hierarchical groups. Neurons were separated from non-neuronal groups based on ABC Atlas taxonomy’s hierarchical clustering which identified 29 neuronal classes and five non-neuronal cell classes. Classification was also available for 15 excitatory-, 12 inhibitory- and two modulatory neuron classes which were easily identified through their main neurotransmitter (Glut, GABA, glycine, dopamine, serotonin), respectively. Hierarchical classification was also helpful with non-neuronal cell types. Astrocytic, oligodendrocytic, and microglial clusters were grouped into a single glia category. Astrocytes were selected from the astro-ependymal cell class, while oligodendrocytes were all clustered into the 31 oligodendrocyte progenitor cells class which contained both mature and precursor oligodendrocytic cell types. All other non-neuronal cell types (i.e. ependymal-, vascular-, other immune cells, olfactory ensheathing cells) were grouped as other non-neuronal cell types (but not part of glial cells). We then used the neuron scaling rules for primate brains [5] to set an upper limit for the three major brain regions: 12,618,420 neurons in the cerebral cortex, 41,825,100 neurons in the cerebellum and 16,446,480 neurons in the rest of the brain. In this way, we preserved the relative differences between regions and cell type groups within the cerebral cortex, cerebellum, and other brain areas, ensuring that the overall numbers aligned with the required values. The only divergence from this occured in the cerebellum, where the ABC Atlas does not distinguish between the granular, molecular, and Purkinje layers within the cerebellar lobes. By subdividing the lobes into their leaf regions, we enabled separate scaling of excitatory neurons in the granular layers (while keeping inhibitory cell type numbers constant) and scaling of non-neuronal cell types across all cerebellar layers.

In the cell type density transplant method we allowed peer-reviewed literature values — specifically on neuron counts — or user provided inputs to serve as ground truth for any number of brain regions. In case literature cell count values could be translated into clusters, we edited the density value of the set of clusters, and transplanted specific density values in the region. Of note, to globally scale the brain and set the final number of neurons to 70.89 million neurons, we took into consideration that some regions received their density through density transplant, and scaling from the first method took effect in only non-transplanted regions, while keeping transplanted densities constant.

We devised a third scaling method, based on the average Nissl template. In this method, we scaled densities to precisely reflect Nissl intensity values in every brain region. The intensity of Nissl staining correlates with cell density, particularly when assessed at a global scale, though variations in cell size and staining intensity introduce non-linearity to this correlation [31,32]. We calculated the average intensity in every brain region in the enhanced average Nissl template [29]. The average Nissl template was created from 86,901 coronal Nissl-stained slices from 734 post-natal day 56 C57BL/6J mouse brains from the AIBS *in situ* hybridization data portal [29]. We proposed two ways to match Nissl intensity values, either by: 1) matching a specific intensity value as the scaling factor (e.g. the highest average Nissl intensity is measured in the crus 2 lobule’s granular layer and we set the density for this intensity to 4 million cells/mm^3^) or 2) assigning the lowest Nissl intensity value, observed in the lateral preoptic area, to correspond to its specific cell density. A scaling factor was then applied based on the difference in Nissl intensities across regions. This way the cumulative densities of cell types matched what the Nissl volume predicted to be a relative density in the given region (Fig. 2S3). Density values were proportional to Nissl intensity values in the average Nissl volume, with total cell counts determined by the initial conditions. Within each region, the relative cell type ratio then specifies the final density of each cell type. However, the correlation between Nissl staining and cell density was not perfectly linear. Nissl staining intensity can vary due to factors like cell size, cell type composition (neurons vs. glia), and RNA content within cells (particularly the ribosomal RNA in the rough endoplasmic reticulum), which means areas of equal cell density might not always exhibit the same staining intensity. Nevertheless, within certain controlled conditions, Nissl intensity was a useful proxy for relative cell density across different brain regions.

### Voxel-Level granularity using Nissl intensity

As cell type densities are calculated for each leaf region, densities remain constant within each region and do not vary on a voxel-by-voxel basis across the brain, except at regional borders. To address this limitation, we used the average Nissl template [29] and calculated voxel-level granularity within each brain region by normalizing Nissl intensity values relative to the mean. These normalized voxel values were then applied to each cell density in the 3D representation. We chose this template due to its alignment with the extended CCFv3 annotation volume, though other Nissl-stained slices can also be used (Fig. 2S1). The benefit of this adjustment was to change the constant average values of any given cell type across a region and better reflect the variability. This step is optional, but one limitation to consider is that each cell type is normalized with the same Nissl-derived factor across its region, even though Nissl intensity represents the cumulative signal of all cell types in a given voxel, regardless of their specific contributions or ratios in that area. This approach maintains the region’s overall density by keeping the cell-type ratios constant; however, we lack information on the true ratio of cell types from one voxel to another. In summary, our pipeline provides the ability of scaling or transplanting densities across the whole brain with voxel level granularity while leveraging the benefits of the transcriptomic atlas’s cell type resolution. At this stage of the pipeline, various types of biologically relevant experimental data can be incorporated into the 3D atlas and converted into transcriptomic densities.

### Alignment of mRNA to established neuronal t-types

One of the first steps of the pipeline was to assign reference t-types defined by Yao et al. in their scRNA-seq study to patch-seq datasets using mRNA data (Fig. 1 B*ii*)[8]. We used the trimmed means (25% - 75%) of Yao et al. scRNAseq dataset as reference points for the t-types alignments. To process and align gene expression profiles, we integrated several libraries, including *scanpy*, *mygene*, and *UMAP* [33,34]. We first cleaned the t-types from Yao et al., to create a reference set of gene expression profiles for unique cell types. We selected only neuronal t-types and duplicate gene identifiers were removed to ensure only unique genes were retained.To analyze patch-seq data, we merged exon and intron counts from single-cell mRNA data and filtered cells based on RNA quality, retaining only high-quality samples. Gene selection was performed based on non-zero mean expression levels [28] , which allowed us to identify a subset of genes for alignment purposes. In brief, we used thresholds for expression frequency and mean log-transformed expression to identify an optimal subset of genes for downstream analysis (i.e. genes with high frequency of zero expression and high level of nonzero expression).

For alignment, we mapped common genes between the Yao reference dataset and the patch-seq dataset using the *ingest* function from the *scanpy* package, standardized the merged datasets, and used *UMAP* to reduce dimensionality. We computed the nearest-neighbor relationships between the reference t-types and patch-seq profiles using principal component analysis (PCA) followed by calculation of the Euclidean distance between the patch-seq and the t-type projections. Subsequently, each patch-seq cell was assigned to its closest reference t-type.

To visualize the alignment results, we generated heatmaps showing the distribution of Patch-seq cells relative to reference t-types. The cells were further divided into excitatory and inhibitory subtypes to enhance interpretability. We used this data to illustrate the alignment fidelity for both excitatory and inhibitory cell populations (Fig. 1S1).

### Morphological classification

Morphological classification was conducted using neurons from various brain regions with data obtained from the Blue Brain dataset [35], combined with Gouwens [27] and Scala [28]. Morphological reconstructions were processed with the *morphology-workflows* package to curate and align the different datasets [36]. Subsequently, morphological features were extracted using the *Topological Morphology Descriptor (TMD)* package [37]. *TMD* provides a topological description of neuron morphologies, capturing key characteristics of neuronal morphologies that can be used for classification and clustering of neurons [38]. Unsupervised clustering was conducted on the vectorized embedding of persistence images [39] of the TMD representation of neurons, applying a customized K-means clustering algorithm which produced approximately 200 distinct morphological types across diverse brain regions. Clustering was performed independently for inhibitory and excitatory neurons, as well as separately for dendritic and axonal structures. This approach yielded 10 clusters per neurite category (dendritic or axonal) and per neuron main type (inhibitory or excitatory).

Canonical m-types were named following a standardized convention to reflect both neuron main type and clustering results, formatted as follows:

NeuronType_DendriticCluster_AxonicCluster (e.g., IN_DEND_0_AX_2 for inhibitory neurons with dendritic cluster 0 and axonic cluster 2). This schema resulted in 200 potential m-types, derived from the combination of 2 neuron types, 10 dendritic clusters, and 10 axonic clusters. However, fewer m-types are observed in practice for the data of Scala and Gouwens, as not all dendritic and axonal cluster combinations are present in the datasets (see supplementary m-type list). Morphologies from patch-seq datasets were then assigned an m-type based on their projected location within the m-feature space.

The probability of observing a specific m-type given a t-type was derived by analyzing their occurrences in the patch-seq datasets. We defined:

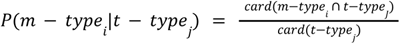

where *card(x)* is the cardinal (i.e. the number of elements) of the set *x*.

### Electrophysiological classification

To compute the conditional probabilities of observing specific e-types based on t-types, we used data from a reference dataset [40] alongside the patch-seq datasets. First, electrophysiological features (e-features) were extracted from recordings using the *EFEL* package, following established protocols [41]. To standardize the data and mitigate variability induced by input resistance, traces were relabeled as a percentage of the rheobase current. For classification, we focused on features extracted from traces presenting rheobase percentage available across all cells. Data cleaning was performed to retain only the shared feature columns, eliminating rows and columns with missing values to ensure consistency across datasets. Each e-feature dataset was subsequently standardized with *StandardScaler* from *scikit-learn (1.4.2)*[42].

To cluster these features, we then used hierarchical clustering (via *AgglomerativeClustering* from *scikit-learn (1.4.2)*) to assign each sample to one of 20 feature-based clusters common to the patch-seq and reference datasets. Selected features were prioritized for their relevance to electrophysiological behavior while maintaining independence from t-type labels, thus capturing essential biological variation across cell types. We used previously published data as a reference dataset that offers a collection of traces with reference labels from the Petilla convention (e.g. cNAC, [40]). The obtained clusters helped structure the data for probabilistic mapping. The probabilities of observing an e-type given a t-type were computing as done previously [41] according to the following equation:

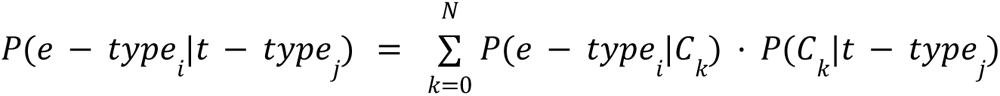

with

- *e* – *type_i_* the ensemble of elements with the *i^th^* e-type in the reference dataset (SSCx).
- *t* – *type_j_* the ensemble of elements with the *j^th^* t-type in the patch-seq datasets.
- *N* the number of common clusters.
- *C_k_* the *k^th^* cluster.
- *N_e-types_* the number of e-types.
- *N_t-types_* the number of t-types.
- 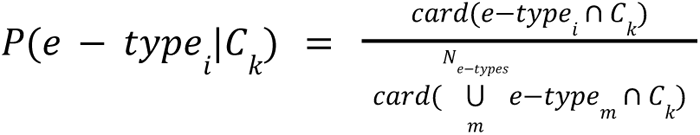
- 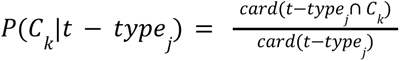

To further analyze t-type and e-type correspondences, a probabilistic mapping matrix was computed to show conditional probabilities for each t-type/e-type pair. This matrix was visualized as a heatmap (Fig. 1S1), illustrating the probability of observing particular e-features given the assigned t-type. Additional post-processing on specific labels was carried out to adjust for excitatory (Glut) and inhibitory (GABA) cell classifications.

### Probabilistic mapping and extrapolation

To compute the joint probabilities of observing combined morphological-electrophysiological types (me-types) given transcriptomic types, we utilized two initial probability matrices: *P*(*m* − *t*|*t* − *type*) , representing the probability of observing a m-type given a t-type, *P*(*e* − *type*|*t* − *type*) representing the probability of observing an e-type given a t-type. We combined these matrices under the assumption that m-type and e-type classifications are independent of each other, allowing us to approximate the joint probability

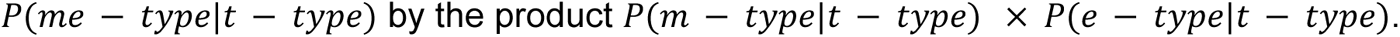

In the implementation, each column of the *P*(*e* − *type*|*t* − *type*) matrix (i.e., each e-type) was used to scale *P*(*m* − *type*|*t* − *type*) element-wise. This was accomplished by iterating through the columns of *P*(*e* − *type*|*t* − *type*), multiplying *P*(*m* − *type*|*t* − *type*) by the e-type probabilities for each t-type. Columns were then renamed to reflect the combined me-type structure (e.g., m-type|e-type). This approach yields a final reduced probabilistic map *P*(*me* − *type*|*t* − *type*) that approximates the joint distribution of me-types for each t-type as a first-order estimate.

To extrapolate me-types for t-types not represented in the patch-seq dataset, we implemented a multi-step approach. Initially, we identified gene markers pertinent to the electrophysiological and morphological properties of neuronal types by applying feature selection methods across various gene expression profiles. RNA-seq data from both covered and uncovered t-types were preprocessed, and feature selection for e-features was conducted using MultiTaskLassoCV and Random Forest regression models, while morphological features (m-types) were analyzed using Logistic Regression with cross-validation, Random Forest classifiers, and mutual information classification from *scikit-learn (1.4.2)*[42].

Top-ranking genes most predictive of m/e-labels were identified based on importance scores derived from each model (e.g. Gini importance, [43,44])(Fig. 1D). A secondary model was then trained to predict the region of origin, and the 100 genes with the lowest scores in this context were identified. The intersection of these two gene sets formed a subspace optimized for predicting m/e-types while minimizing regional variability. The selected genes served as input to a k-nearest neighbors model (Fig. 1E, *i*), where gene expression profiles were scaled, and Euclidean distance matrices were computed to identify the closest matching neuronal types. Probabilistic mappings for uncovered t-types were inferred by calculating the average of the nearest neighbors’ probabilities, weighted by the distance metrics (Fig. 1E,*ii*).

Ten-fold cross-validation techniques (KFold and StratifiedKFold) were employed to assess the performance of different parameter choices and error metrics, including mean squared error (MSE) and mean absolute error (MAE), which allowed us to identify the optimal number of neighbors (best *N_neibhbors’_*, Fig. 1S1). Best results were obtained with the Random Forest algorithms and with *N_games_* = 100 and *N_neibhbors’_* = 5 This extended probabilistic map, integrating the newly extrapolated t-types, was used to later compute the me-type densities across the brain using

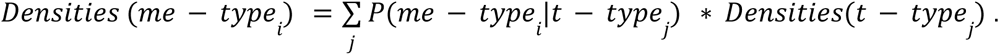

### Calculation of mRNA densities

For verification (see Results, Fig4. B and C), we computed mRNA densities based on either t-type or me-type densities. Specifically, mRNA densities from t-types were calculated as follows:

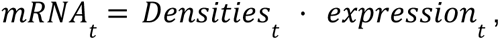

where *expression_t_* is a matrix of mRNA counts (with t-types as rows and genes as columns), *Densities_t_* is a matrix of t-type densities (with brain regions as rows and t-types as columns), and *mRNA_t_* is a matrix of mRNA densities (with brain regions as rows and genes as columns).

Similarly, mRNA densities can be derived from me-type densities as:

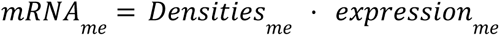

which can be expanded to:

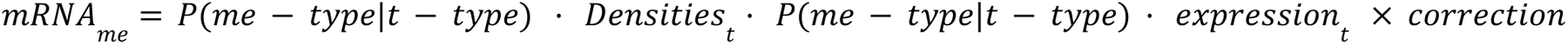

Here, *P*(*me* − *type*|*t* − *type*) represents the previously computed extended probabilistic map, and *correction* is an adjustment factor. This correction addresses the increase in density values that arises when converting 4,804 neuronal t-types into 458 functional me-types, which would otherwise lead to an overestimation of mRNA densities in me-types. The correction factor is defined as:

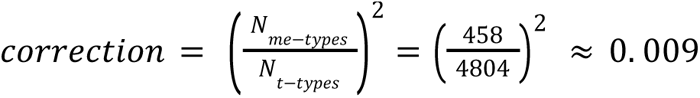

The square term results from the probabilistic map being applied twice in this calculation.

The error between both estimates was computed as 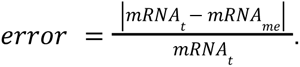.

### Elbow method to determine the number of clusters

To analyze the diversity of cell-type compositions across brain regions (categorized by either t-types or me-types), we performed K-means clustering on UMAP-projected cell-type density data (Fig. 5 A*i*, *ii*). The optimal number of clusters was determined using the Elbow Method, a heuristic that balances cluster compactness with model complexity [45,46]. This approach involves calculating the within-cluster sum of squares (WCSS) over a range of cluster counts, from 1 up to a predetermined maximum (in this case, 10 clusters). WCSS, which measures the total variance within each cluster, serves as an indicator of clustering compactness as clusters are added. When WCSS values are plotted against the number of clusters, the optimal cluster count is identified at the “elbow point,” where additional clusters yield diminishing reductions in WCSS. This elbow point represents a balance between minimizing intra-cluster variance and avoiding unnecessary model complexity.

## Results

To work out a widely usable density atlas we adapted the parcellation annotation with 4-5 levels of hierarchy. To make it compatible with the original brain hierarchy used by older versions of the anatomical annotation volumes, we introduced missing leaf regions as a substructure. We handled gray and white matter regions in a similar manner. In the case of gray matter, we split those substructures which could be further split into leaf regions (the term leaf represents the deepest level of the hierarchical ontology). For white matter, we initially treated parent regions homogeneous without dividing them into their leaf regions. Later, their leaf regions were assigned the same density information as their parent regions. This gave us the flexibility to select a brain region at any level of the hierarchy, if needed, by concatenating its substructures.

Processing all MERFISH slices yielded 5,274 individual 3D brain volumes for each cell type cluster within the ABC Atlas (Fig. 1 A*iii*). Additionally, we grouped cell types based on taxonomic hierarchy, creating 3D density volumes for major cell groups such as excitatory neurons, inhibitory neurons, modulatory neurons, glial cells, and all non-neuronal cells, among others (Fig. 2). In each brain volume, leaf regions contain the estimated average cell density of the specified cell type, calculated from MERFISH slices that intersected the region. For normalization and validation, we also generated brain volumes where voxels represent total cell counts in each region for the major cell groups. To reduce file numbers, we calculated cumulative densities of major cell groups at each level of the anatomical hierarchy, allowing them to be stored efficiently in a single table.

The volumetric region data calculated for each individual cell enabled us to estimate the total cell count or cell type density across the brain by summing the cells of each type within a given region and dividing by its regional volume. This also enabled us to estimate cell densities, which could then be compared to values from the literature (Fig. 3). One of the most reliable measurements of the mouse brain [5] estimated the total number of cells to be 108.69 ± 16.25 million, with the total neuron count at approximately 70.89 ± 10.41 million. Thus, using the estimates of 17.8 ± 3.4% neurons in the cerebral cortex and 59.0 ± 5.0% in the cerebellar cortex, we can calculate that there are approximately 12,618,420 neurons in the cerebellar cortex, 41,825,100 neurons in the cerebellum, and an estimated 16,446,480 neurons in the remaining regions of the brain.

**Figure 3:**
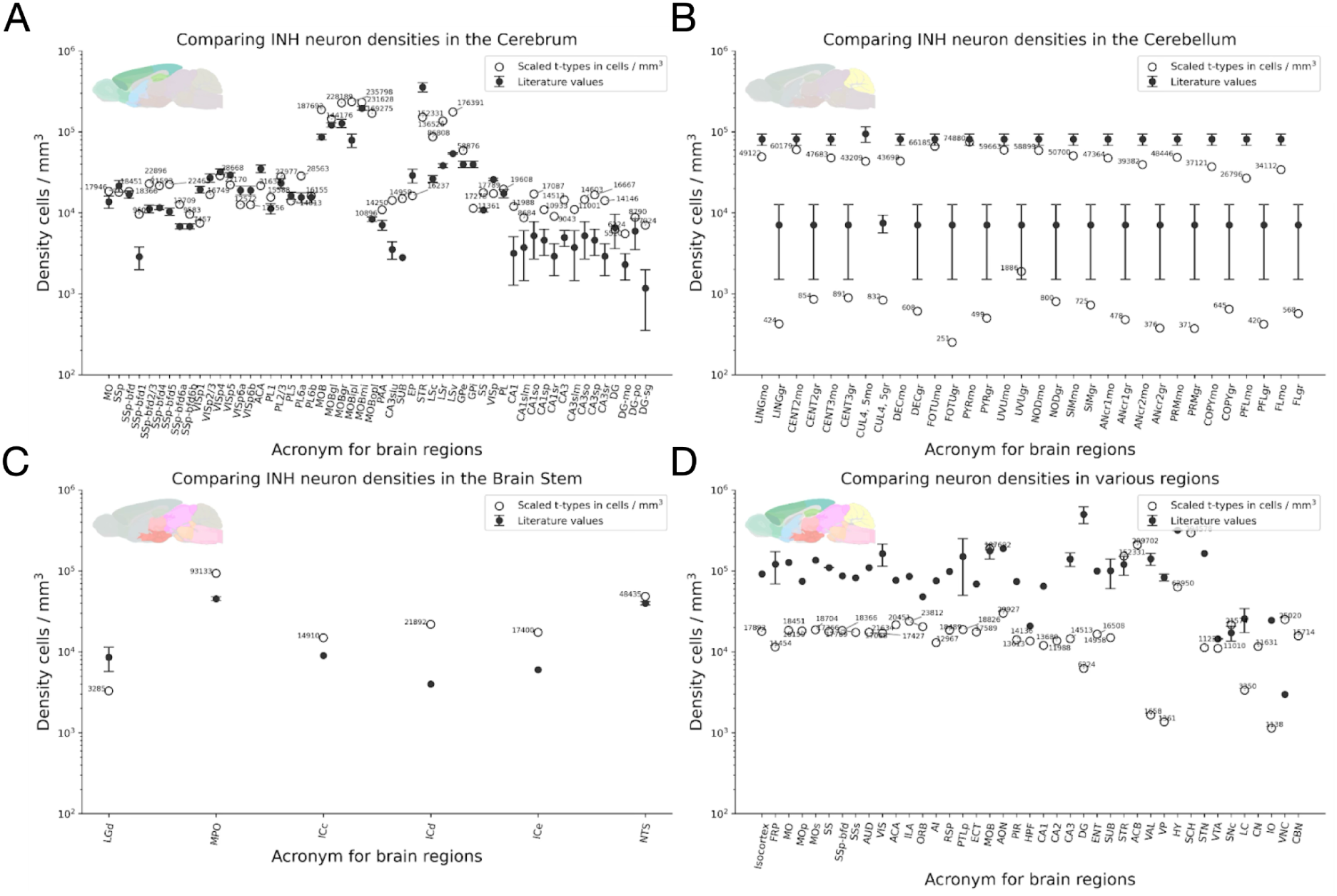
Comparison of predicted density with literature values. **A.** We extracted inhibitory neuron densities from the Cerebrum from the unscaled atlas and compared them with inhibitory neuron numbers from the literature. **B.** Extracted from the Cerebellum **C.** Extracted from the brain stem **D.** Comparing total neuron density values from the unscaled atlas with values from the literature [7].

Since the slices are aligned to the common coordinate framework, the voxelized volume can be used to obtain the total cell and neuron counts. Consequently, MERFISH slices indicate a total of 92 million cells across the entire brain. A surprising finding is that, despite the cerebellum being significantly underrepresented with only 6.2 million cells and 3.5 million neurons, the rest of the brain contains many more cells than anticipated. Our conclusion is that MERFISH slices can not fully capture the true density in highly cell-dense regions, such as the granular layers of the cerebellum or the dentate gyrus of the hippocampus. This discrepancy could be attributed to technical limitations encountered in areas of densely populated cellular regions and/or small cell size making it difficult to define clear cell boundaries. Conversely, the overestimation of cell numbers in other regions, particularly in the fiber tracts and cerebral nuclei, can not be explained by these technical limitations. In our estimates, compared to the neuron scaling rules for primate brains [5], the cerebral cortex has 2.5 times more neurons, while the isocortex contains 1.5 times more. Current understanding considers the neuron count in the fiber tracts to be negligible. Surprisingly, the ABC Atlas identified 61% of all cell types present in the fiber tracts, making this region even richer in cell types than the entire cerebral cortex. This holds true even though, based on the raw MERFISH counts, only 8.81% of the cells are neurons, while 91.18% are non-neuronal. Since we found no significant differences in the QC metrics (provided by the AIBS) between this region and other regions, or between neurons and other cell types within this region, we chose to retain them in the final count, recognizing their potential relevance for capturing important biological information. The estimated total cell numbers in the hippocampal formation align with existing literature, but this is primarily due to the low rate of cell discovery in the densely populated pyramidal and granule cell layers. The thalamus fell within a similar range, while the striatum had more than twice the number of neurons and cells reported in earlier studies. Notably, there was a significant discrepancy in the olfactory areas, which had an estimated 7 million neurons compared to the expected 1-2 million. Similarly high cell counts were found in the cerebral nuclei. It was particularly surprising that low-density areas are consistently overestimated compared to estimates derived from isotropic fractionation. See **Table 1** for more details.

**Table 1:**
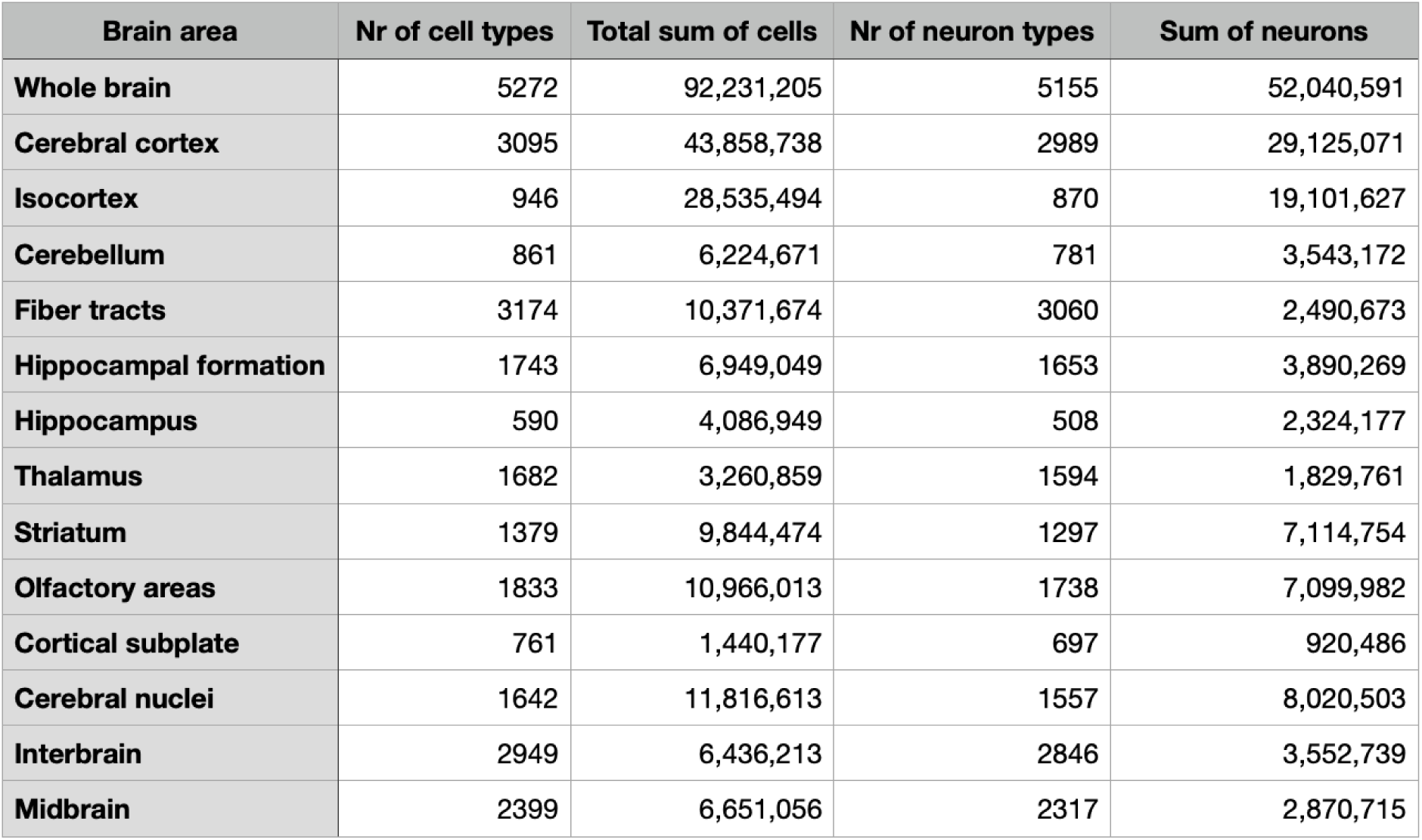
Total cell and neuron counts across the major brain regions, as estimated from MERFISH data. We aggregated the density values and transcriptomic cell type counts across major brain areas, expressed as the number of cells/mm³.

| Brain area | Nr of cell types | Total sum of cells | Nr of neuron types | Sum of neurons |
| --- | --- | --- | --- | --- |
| Whole brain | 5272 | 92,231,205 | 5155 | 52,040,591 |
| Cerebral cortex | 3095 | 43,858,738 | 2989 | 29,125,071 |
| Isocortex | 946 | 28,535,494 | 870 | 19,101,627 |
| Cerebellum | 861 | 6,224,671 | 781 | 3,543,172 |
| Fiber tracts | 3174 | 10,371,674 | 3060 | 2,490,673 |
| Hippocampal formation | 1743 | 6,949,049 | 1653 | 3,890,269 |
| Hippocampus | 590 | 4,086,949 | 508 | 2,324,177 |
| Thalamus | 1682 | 3,260,859 | 1594 | 1,829,761 |
| Striatum | 1379 | 9,844,474 | 1297 | 7,114,754 |
| Olfactory areas | 1833 | 10,966,013 | 1738 | 7,099,982 |
| Cortical subplate | 761 | 1,440,177 | 697 | 920,486 |
| Cerebral nuclei | 1642 | 11,816,613 | 1557 | 8,020,503 |
| Interbrain | 2949 | 6,436,213 | 2846 | 3,552,739 |
| Midbrain | 2399 | 6,651,056 | 2317 | 2,870,715 |

To address the limitations of MERFISH-based estimations, we could enhance its accuracy by scaling the density values (Fig. 3S). However, the challenge with scaling was that it relies on the adoption of a scientifically reviewed experiment from the literature as the reference point. Therefore, we proposed offering multiple methods for scaling density values across the brain to accommodate different use cases and expectations. Of note, the true value of this transcriptomically defined cell type dataset lies in its comprehensive collection of cell density values across both large and small brain regions, all gathered using the same technology. This consistency allowed us to avoid the increased variance, cross-platform bias, and batch effects. Although the total cell numbers may be skewed due to the technological constraints and extrapolation, the ratios of cell types, as well as their presence or absence, provided highly valuable insights. Taking these factors into account, we recommended these adjustments to the total density values of cell types while preserving their ratios across regions:

The first scaling method involves aligning the global values with those obtained through isotropic fractionation, specifically the neuron scaling rules for primate brains. This approach allowed us to scale the total counts of neurons to 70.89 million neurons and the total cell counts (glia included) to 108.69 million cells across every brain region. This scaling approach allowed us to adjust cell numbers across large brain regions while preserving the relative densities between regions and cell types. However, a limitation is that certain regions may retain errors and stand out as outliers due to issues such as poor coverage, low-quality slices, or compromised image quality.

In the second scaling technique we wanted to introduce density transplantation. For example, specific modeling studies require a defined set of neuron types, including particular density and standard deviation, often due to computational limitations or unpublished measurements. In our scaling pipeline we provided the ability to swap out the estimates from MERFISH values with specific values for any given cell type or cell class (or at any level of the cell type hierarchy) while keeping the other regions constant. In this case, we can still apply the first scaling method while the pipeline will take into consideration that transplanted regions have to stay constant in any subsequent changes. Since in the first step we need to scale up or down the total number of cells across regions to meet a maximum number (e.g. 70.89 million neurons or 108.69 million with glia included) the neighboring regions in the brain area will get scaled more or less depending on the transplanted values. In this study we transplanted 51 regions (including larger regions e.g. Striatum) where we had estimates of total neuron or inhibitory and/or excitatory cell densities [47]. After transplantation we ran our first global scaling method based on the neuron scaling rules for primate brains while keeping those densities constant which were fixed by the transplant operation.

The third method, average Nissl volume based scaling, also provided a way to scale density values without changing the ratio of individual cell types (Fig. 2S3). It also enabled us to estimate cell densities in regions that were not covered by the MERFISH slices. When comparing the two types of Nissl scaling, we consistently observed that the maximum-based method produced higher cell densities across all brain regions. Minimal differences were noted in the cerebral nuclei, main olfactory bulb, and isocortex, while the hippocampal formation showed similar cell counts except in specific areas where cell diameter is smaller (field CA1, pyramidal layer, field CA2, pyramidal layer, field CA3, pyramidal layer, dentate gyrus, granule cell layer). The most significant differences were found in the cerebellum and certain regions of the cerebral cortex. Overall, unscaled and scaled values exhibited a very strong correlation.

### Probabilistic mapping

#### Pipeline Description

The probabilistic mapping approach is centered around estimating the likelihood of observing specific me-types given a known t-type, based on a multimodal dataset, as previously outlined by [41]. The process begins by labeling the patch-seq dataset with the relevant cell-type classifications. Specifically, we utilized t-types from Yao et al.,e-types following the Petilla convention, and canonical m-types as defined by Kanari et al. (unpublished) (Fig. 1B). To extend this labeling beyond the scope of the patch-seq dataset, we extrapolated probabilities to t-types not represented in this dataset (Fig. 1C, D and E).

#### Probabilistic map pipeline validation and optimization

To assess the robustness of our reduced probabilistic map *P*(*me*∣*t*), we conducted a series of intermediate validations on the alignment methods. First, we used native t-type labels from the patch-seq dataset to compare with assigned reference t-types from Yao et al. (Fig. 1S1). The results were satisfactory, as excitatory neurons from the patch-seq dataset predominantly mapped to cortical t-types (Fig. 1S1, top), while inhibitory neurons retained their major family classifications, such as Pvalb, Vip, or Sst (Fig. 1S1, top). For e-type mapping validation, we calculated the probabilities of observing a t-type given a canonical e-type (Fig. 1S2A). The alignment preserved functional consistency, with excitatory e-types mapping to excitatory t-types and inhibitory e-types demonstrating a similar trend. Additionally, the inhibitory mapping corresponded well with previously documented mappings (Roussel et al., 2022), with e-types like cSTUT, dNAC, and dSTUT showing a preference for Pvalb cells, indicative of fast-spiking profiles.

Although quantitative data was unavailable for morphological validation, we qualitatively assessed the morphological space coverage of the reference dataset. Both patch-seq datasets spanned over 75% of the reference dataset morphological space, indicating substantial overlap (Fig. 1S2B). Taken together, these alignment validations demonstrate that the methods used were sufficiently precise to construct the reduced *P*(*me*∣*t*).

To extend the probabilistic map to uncovered t-types, we conducted a 10-fold cross-validation to benchmark multiple machine learning techniques for feature selection, including random forest, Lasso, and mutual information [42]. This process also optimized the number of genes (*N_genes_*) used to define the reduced space, as well as the number of neighbors (*N_neighbors_*) required for calculating the probabilities of observing me-types for the uncovered t-types. This optimization step yielded the best results using the random forest algorithm with *N_genes_* = 100 and 10 (Fig. 1S3).

#### Me-types densities and gene expression verification

The primary motivation for computing the probabilistic map *P*(*me* − *types*∣*t* − *types*) was to apply it to the t-type densities (generated in the first stage of the pipeline Fig. 1A) according to the equation:

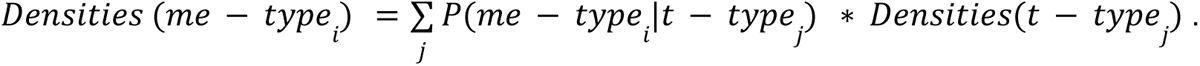

This calculation yields density distributions across all brain regions for the 453 me-types, with examples illustrated in Fig. 4A and 4B. Given the reduction from over 4804 t-types to 458 me-types, some information loss due to noise is anticipated in terms of gene expression fidelity. To quantify this noise, we compared the mRNA counts across brain regions obtained from the t-type densities (Fig. 4B, left) with those derived from the me-type densities (Fig. 4B, middle) by calculating their relative error (Fig. 4B, right, see methods). This comparison reveals that mRNA counts are more uniform across brain regions for me-types than for t-types (Fig. 4B, left and middle). Apart from a few outliers (e.g., MMp), the relative error remains generally consistent across brain regions for individual genes (Fig. 4B, right). As many t-types are mapped to a reduced number of me-types, this increase in homogeneity is expected (see discussion).

**Figure 4:**
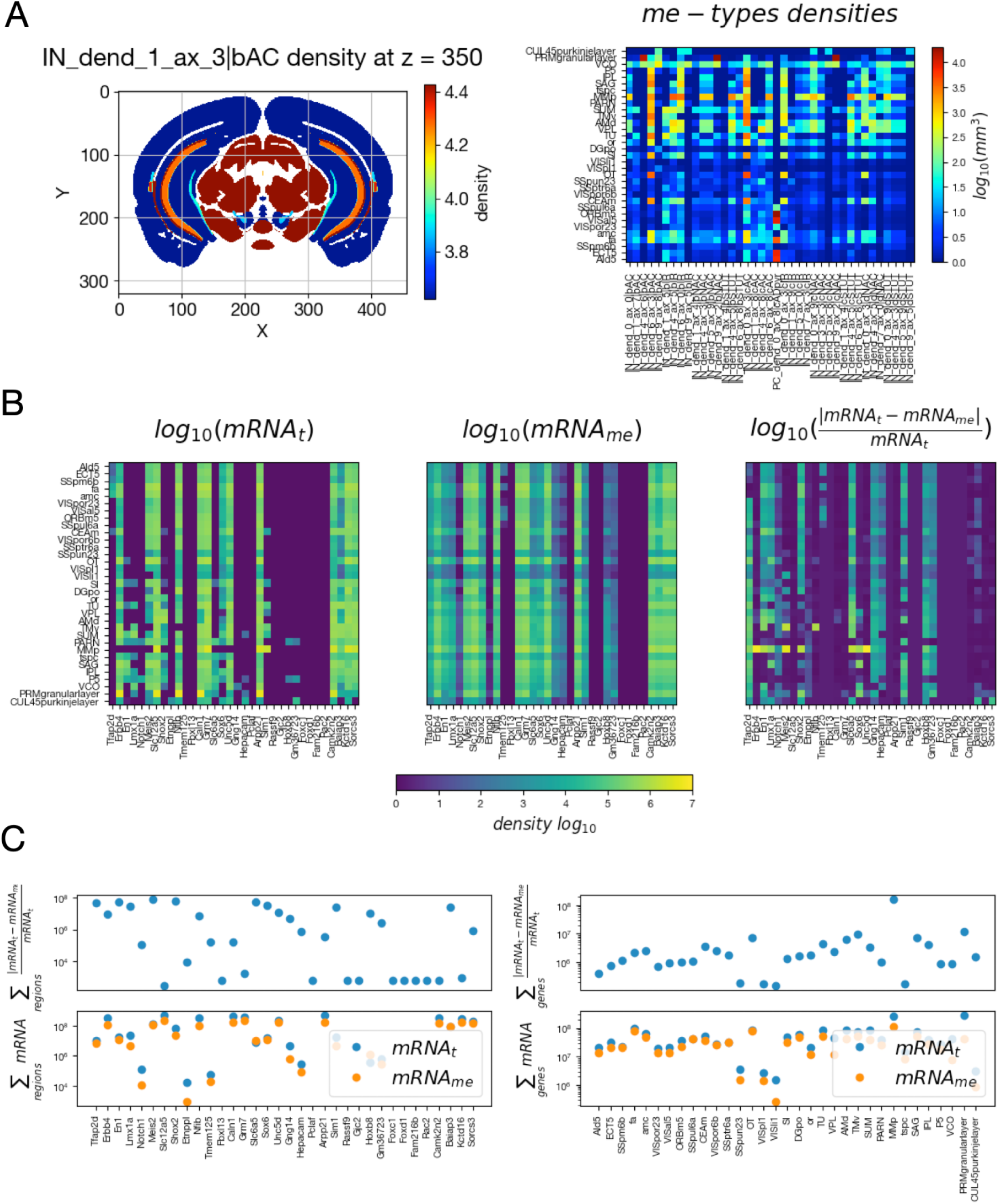
Probabilistic mapping application on t-type densities and validation of mRNA counts globally. **A.** Example of a coronal section of the IN_dend_1_ax_3|bAC me-type density after applying the probabilistic mapping (*left*) and a heatmap (*right*) of obtained densities for a selection of brain regions (rows) and a selection of me-type (columns). Only a selection of brain regions and t-types are shown for clarity. **B.** mRNA counts obtained by multiplying cell types gene expression profiles with their densities (see methods) in a selection brain regions and a selection of genes for t-types (*left*) and me-types (*middle*) and their relative error (*right*). **C.** Semilog plots of mRNA counts summed over regions (*left*) or genes (*right*). The top row shows the relative error while the bottom row shows the summed mRNA counts for t-types in blue and me-types in orange.

**Figure 5:**
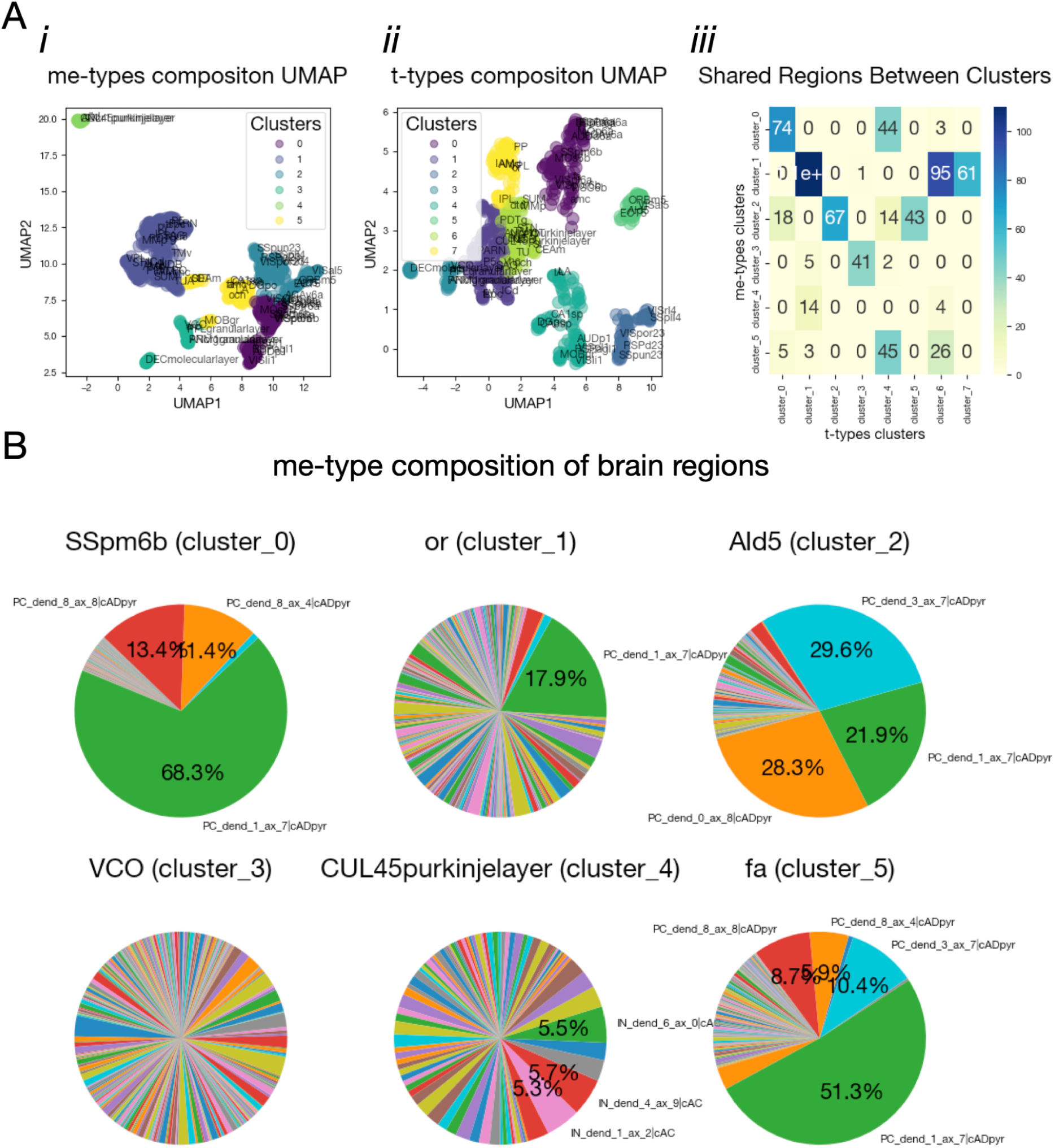
Brain region cell types composition. **A.** UMAP projection of brain region cell type composition when considering me-types (*i*) and me-types (*ii*). Points are color coded according to the *kNN* clustering results. (*iii*) Matrix showing the shared brain region between me-types composition clusters (rows) and t-types composition clusters (columns) **B.** me-types composition of 6 brain regions. Each brain region comes from a different cluster. Only the me-types composing more than 5 % of the brain region are shown.

To gain a more general overview of the differences in mRNA counts between t-types and me-types, we plotted the summed counts on a semilog scale, both across brain regions (Fig. 4C, bottom-left) and across genes (Fig. 4C, bottom-right). We also visualized the summed relative errors (Fig. 4C, top-left and right). Overall, the cumulative mRNA counts across genes and brain regions remained of similar magnitude for both me-types and t-types.

Nonetheless, because the same amount of mRNA has to be split in either 4804 t-types or 453 me-types, we have mechanically more mRNA per me-types than per t-types. Hence, when we look at the cumulative errors for individual genes (Fig. 4C, left), they often exceed those for brain regions (Fig. 4C, right). Meaning that, even though we have more genes than brain regions, when we sum over brain regions, the errors often rise up higher when compared to summing over genes. These results suggest that, with respect to mRNA counts, the error introduced by homogenizing the composition across brain regions is more important than the error introduced by having more mRNA per cell type (i.e. decreasing the number of cell-types from t-types to me-types).

#### Comparison of me-types and t-type composition of brain regions

To evaluate the impact of homogenizing cell-type compositions across brain regions, we clustered brain regions based on their cell-type compositions for both me-types and t-types (Fig. 5Ai, *ii*). On these plots one data point represents a brain region.If two brain regions are located close together, they share similar composition in terms of me-types (Fig. 5 A*i*) or t-types (Fig. 5 A*ii*). Using the elbow method for kNN clustering (see Methods), we identified six clusters for the me-type compositions of brain regions, compared to eight clusters for t-types. While a clear elbow point indicated optimal clustering for me-types, the t-types did not show a distinct elbow, suggesting that a higher number of clusters might be considered for t-type compositions. This observation further indicates that cell-type compositions become more homogenized across brain regions when transitioning from t-types to me-types. However, the relatively small difference in cluster numbers between me-type and t-type compositions implies that the homogenization effect is low.

To determine the consistency between me-type and t-type composition clusters, we examined the extent to which brain regions were shared between clusters derived from me-types composition and those from t-types composition (Fig. 5Aiii). Given the reduced number of me-types compared to t-types, we anticipated that certain clusters of me-type compositions would correspond to multiple clusters of t-type compositions (e.g., me-type clusters 0, 1, and 2; Fig. 5Aiii). Interestingly, we also observed the reverse scenario, where certain clusters of t-types (clusters 4 and 6) mapped to multiple me-type clusters. This bidirectional mapping suggests a potential reduction in the specificity of brain region cell-type compositions when moving from t-types to me-types.

To illustrate these findings, we plotted the me-type compositions of six brain regions, representing one example from each me-type composition cluster (Fig. 5B). The results highlight that some regions are dominated by a few excitatory types (e.g., SSp-m6b or Ald5), while others exhibit more diversity (e.g., VCO). This variation may be explained by the structural organization of cortical regions, which are layer-specific (e.g., layer 6b of the somatosensory primary cortex) and are populated by distinct excitatory me-types (e.g., PC_dend_1_ax_7|CADpyr, which is primarily found in cortical layers 5/6)[11,27,28].

---

**Figure 1 Supplementary 1:**
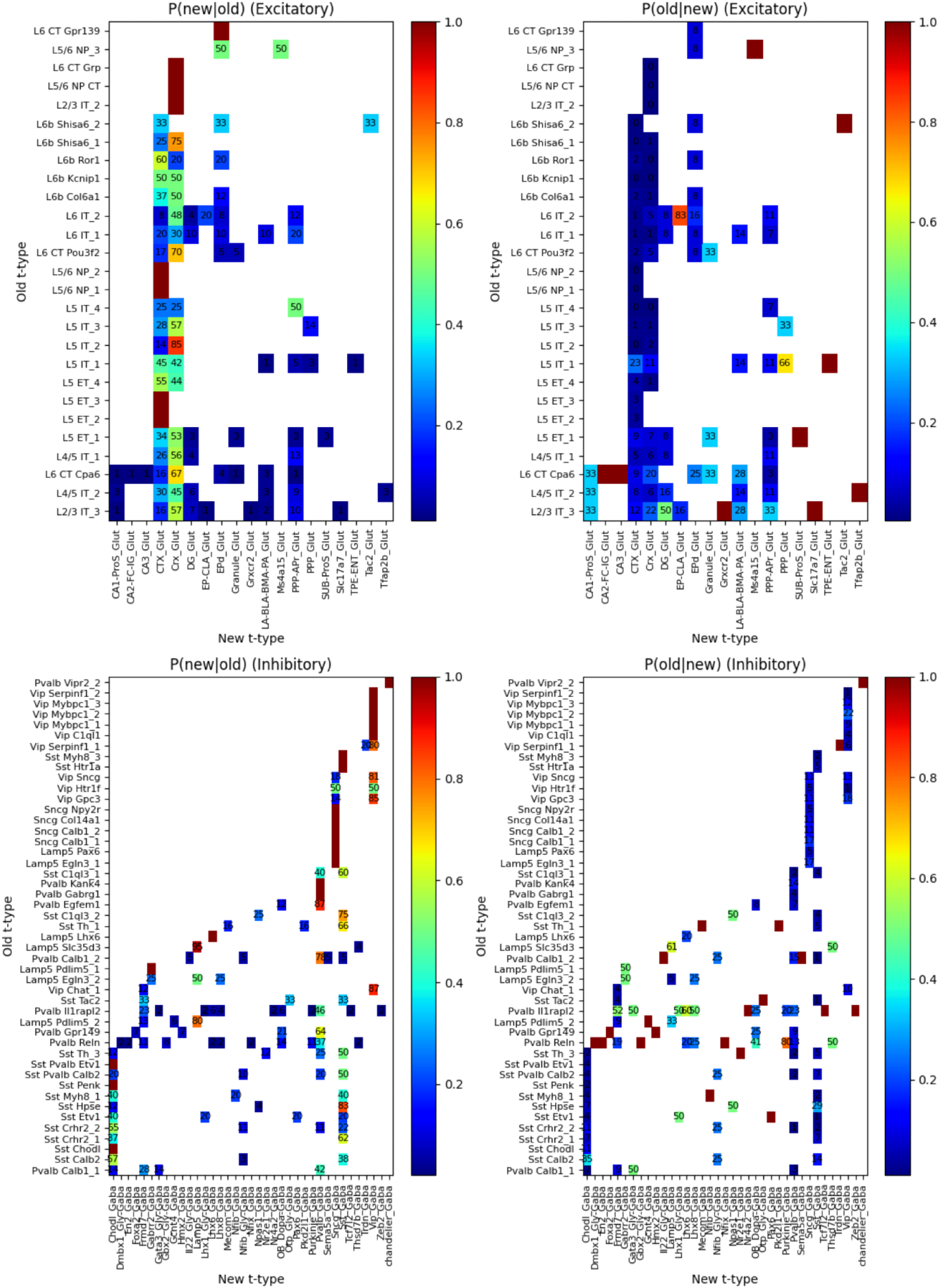
Validation of t-types alignment method. Probability maps highlighting the t-type alignment between native labels (given by the patch-seq dataset) with the assigned reference t-types (yao et al.). Excitatory t-types on top row and inhibitory on the bottom. Probabilities of having a yao t-type given a native label on left, Probabilities of observing a native label given a yao t-type on the right.

**Figure 1 Supplementary 2:**
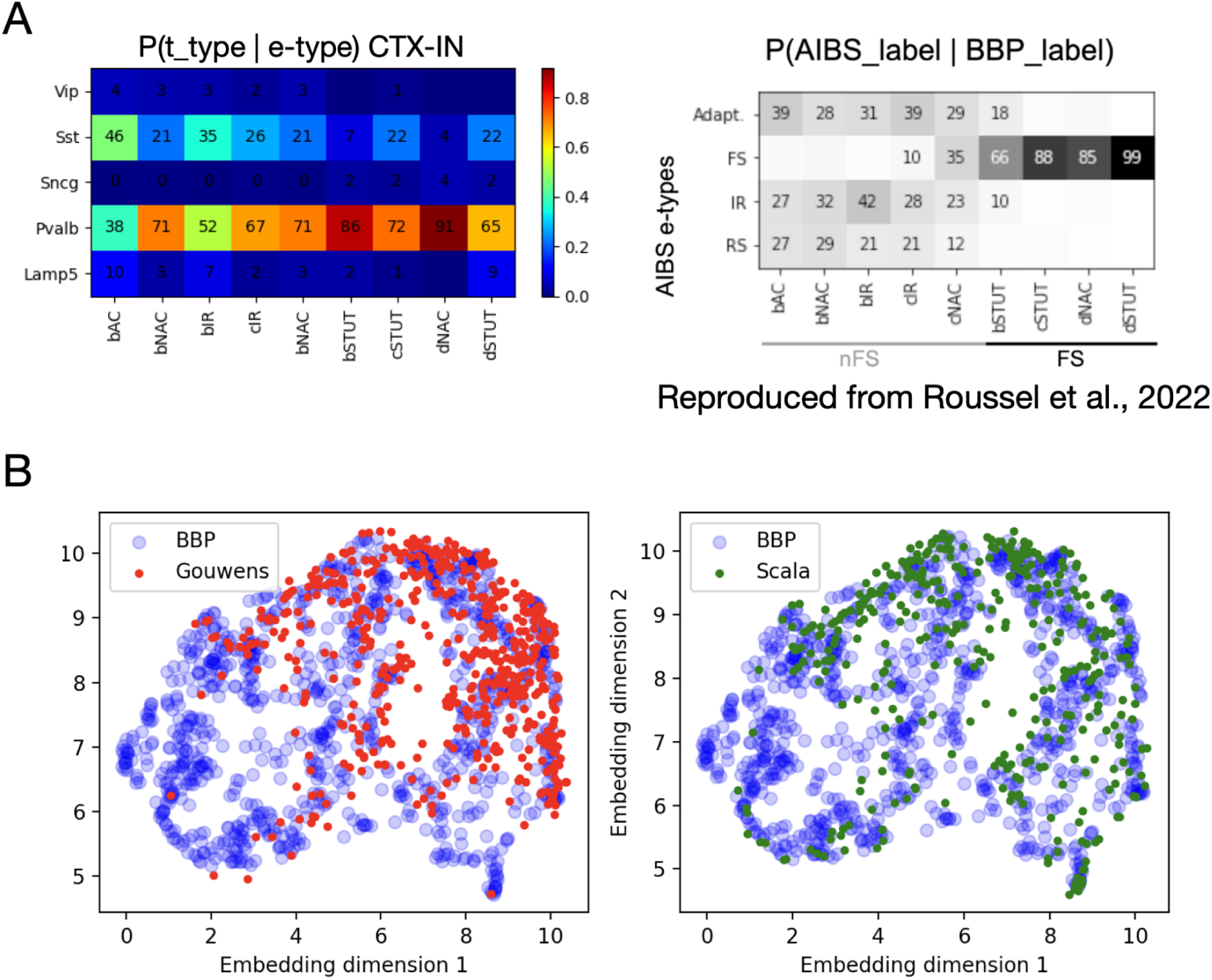
Validation of electrophysiological and morphological mappings. **A.** Probabilities of observing a canonical e-type given a yao et al. t-type in the Scala patch-seq dataset (left). Probabilities of observing e-types as defined by Gouwens et al. 2019 given canonical e-types (right, reproduced from [41]). bSTUT, cSTUT, dNAC and dSTUT mapped preferentially to fast spiking cells (FS) while all the other mapped mostly to non fast spiking neurons (nFS) **B.** Coverage of the Gouwens et al. (2020) dataset (left) and the Scala et al. (2020) dataset of the embedding of the canonical morphological space defined by the reference morphological dataset (labeled as “BBP”).

**Figure 1 Supplementary 3:**
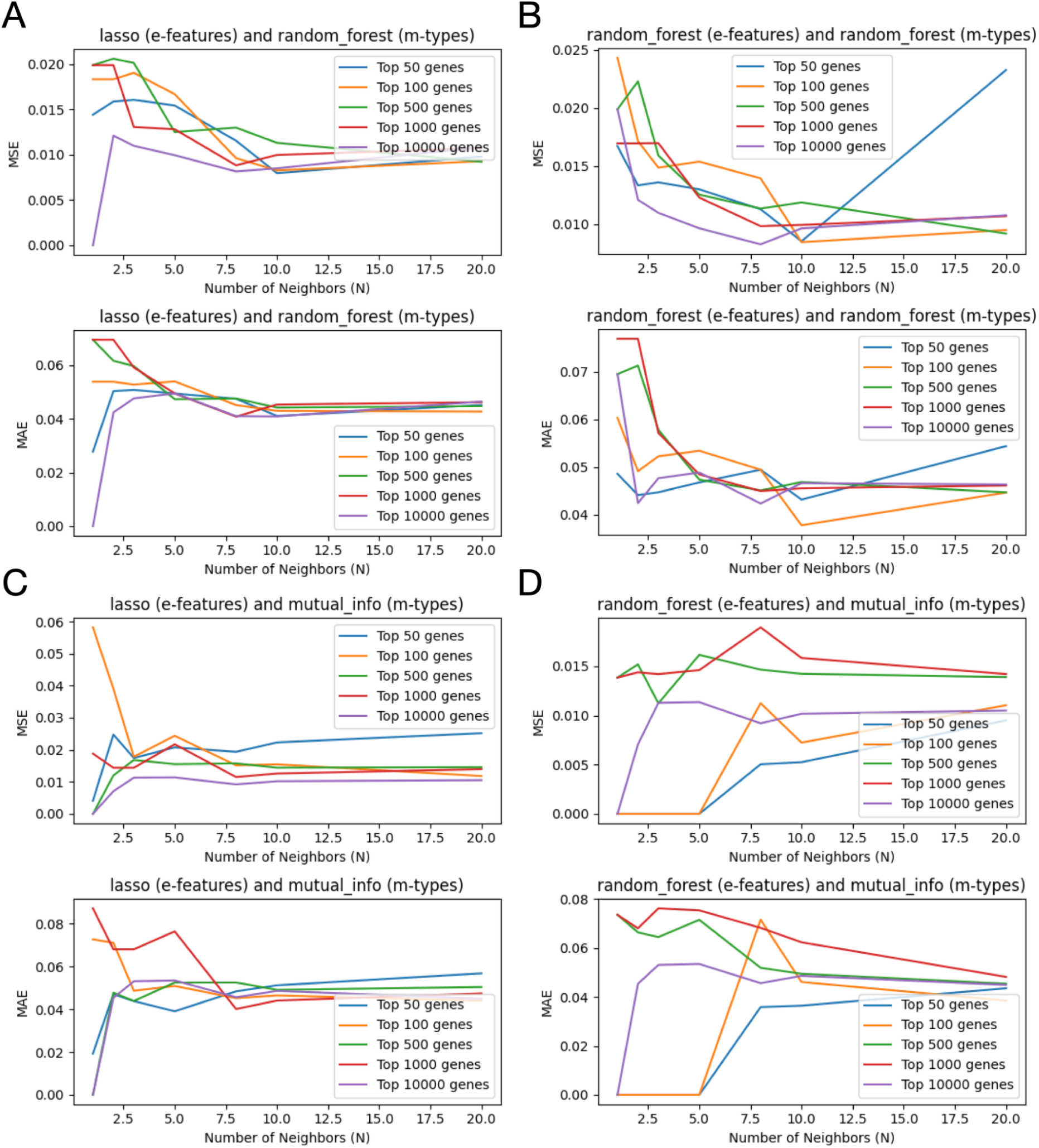
Optimization of the pipeline parameters (*N_genes_*, *N_neighbors_* Clustering algorithm). Mean squared error (MSE, top) and mean absolute error (MAE, bottom) as a function of *N_neighbors_* for *N_genes_* = 50, 100, 500, 1000 *and* 10000 and for multiple machine learning algorithms combination for e-features space and m-types space gene selection for the 10-fold cross-validation process. Several combinations were tried: lasso and random forest (**A**), random forest and random forest (**B**), lasso and mutual info (**C**), random forest and mutual info (**D**).

**Figure 2 Supplementary 1:**
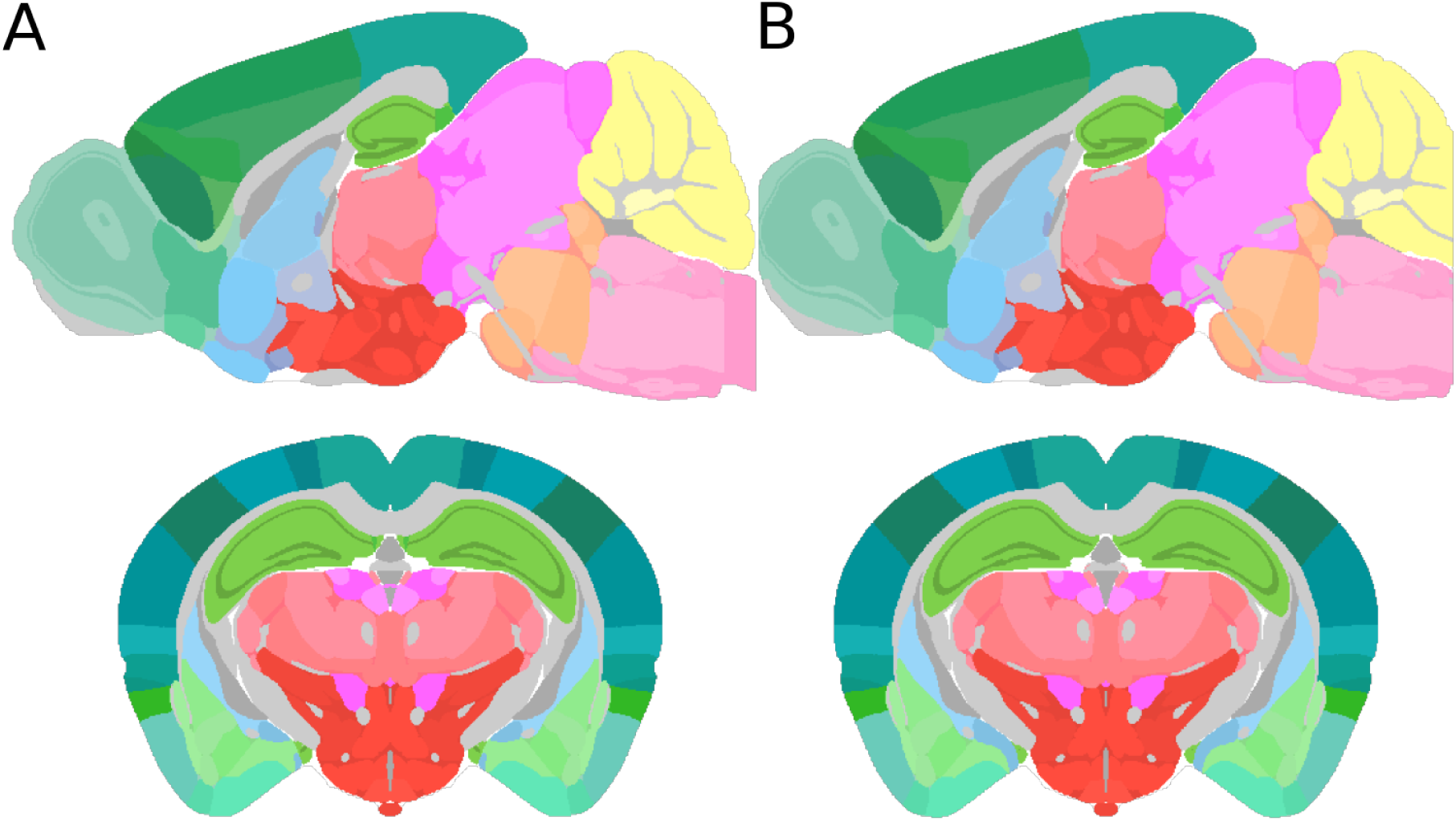
Different annotation atlas versions can be used to create various iterations of transcriptomic atlases. **A.** Sagittal (y=200) and coronal (x=315) sections of the CCFv3 annotation volume (from the AIBS (x=300) and the **B.** extended CCFv3 annotation volume from the literature [29].

**Figure 2 Supplementary 2:**
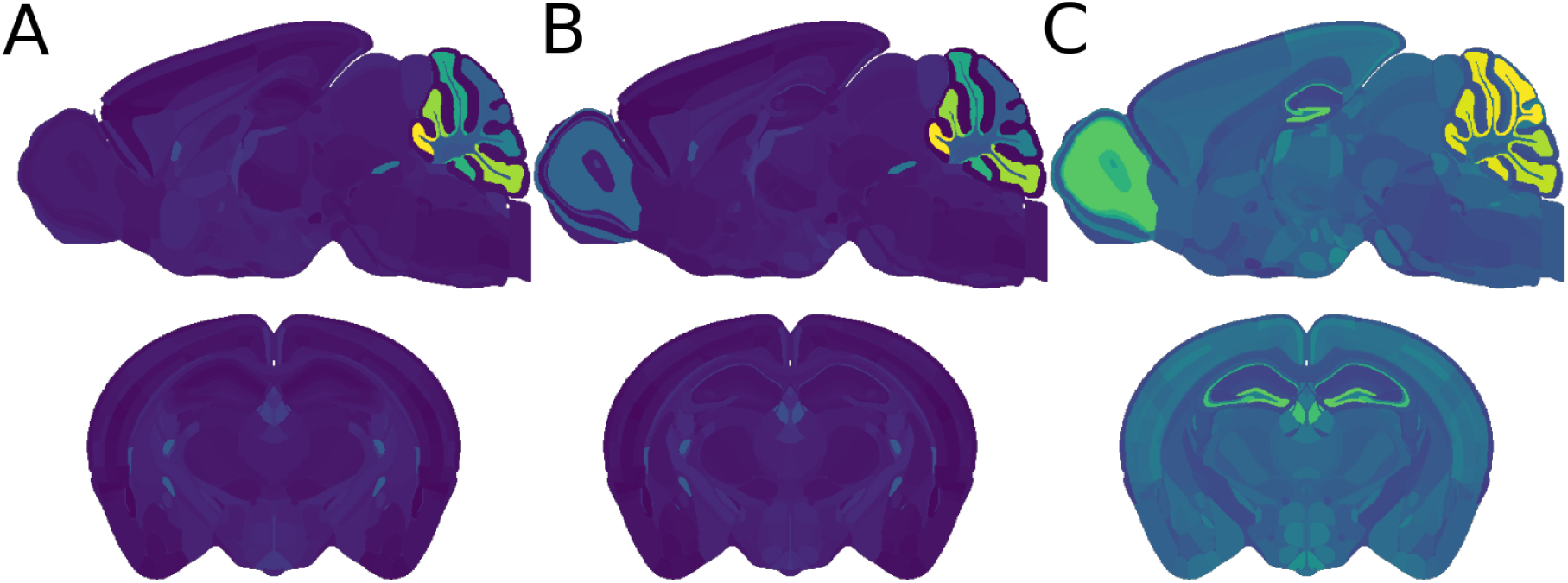
3D transcriptomic atlases generated with scaled densities. Sagittal (y=200) and coronal (x=300) sections of scaled density volumes (every cell types included): **A.** scaled to meet total cell numbers in [5], **B.** scaled to meet total cell numbers in [5] with transplant, **C.** all regional total cell type densities scaled match differences in Nissl intensity.

**Figure 2 Supplementary 2:**
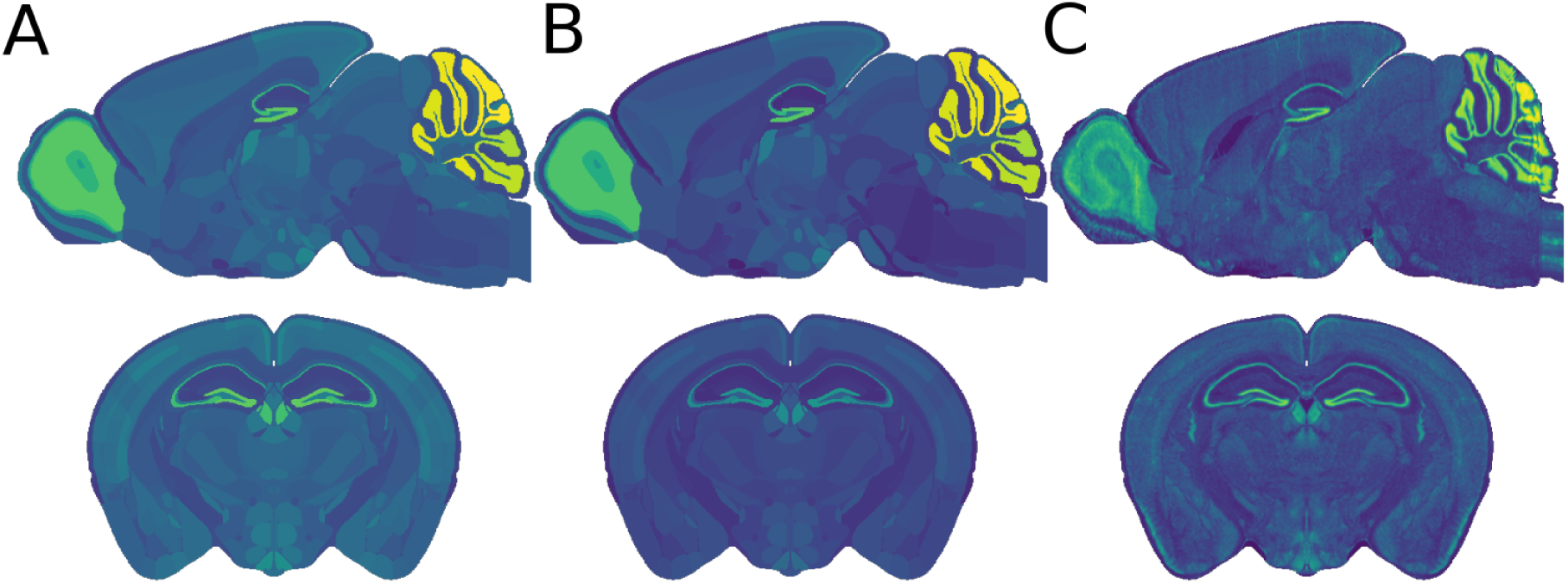
Nissl scaling of 3D transcriptomic atlases Sagittal (y=200) and coronal (x=300) sections of scaled density volumes (total cell): **A.** all regional total cell type densities scaled match differences in Nissl intensity, scaled with the minimum Nissl intensity, **B.** all regional total cell type densities scaled match differences in Nissl intensity, scaled with the Maximum Nissl intensity equals 4 million cells/mm^3^ **C.** Nissl granularity was added to **A** to emulate variance within each region.

**Figure 3 Supplementary 1:**
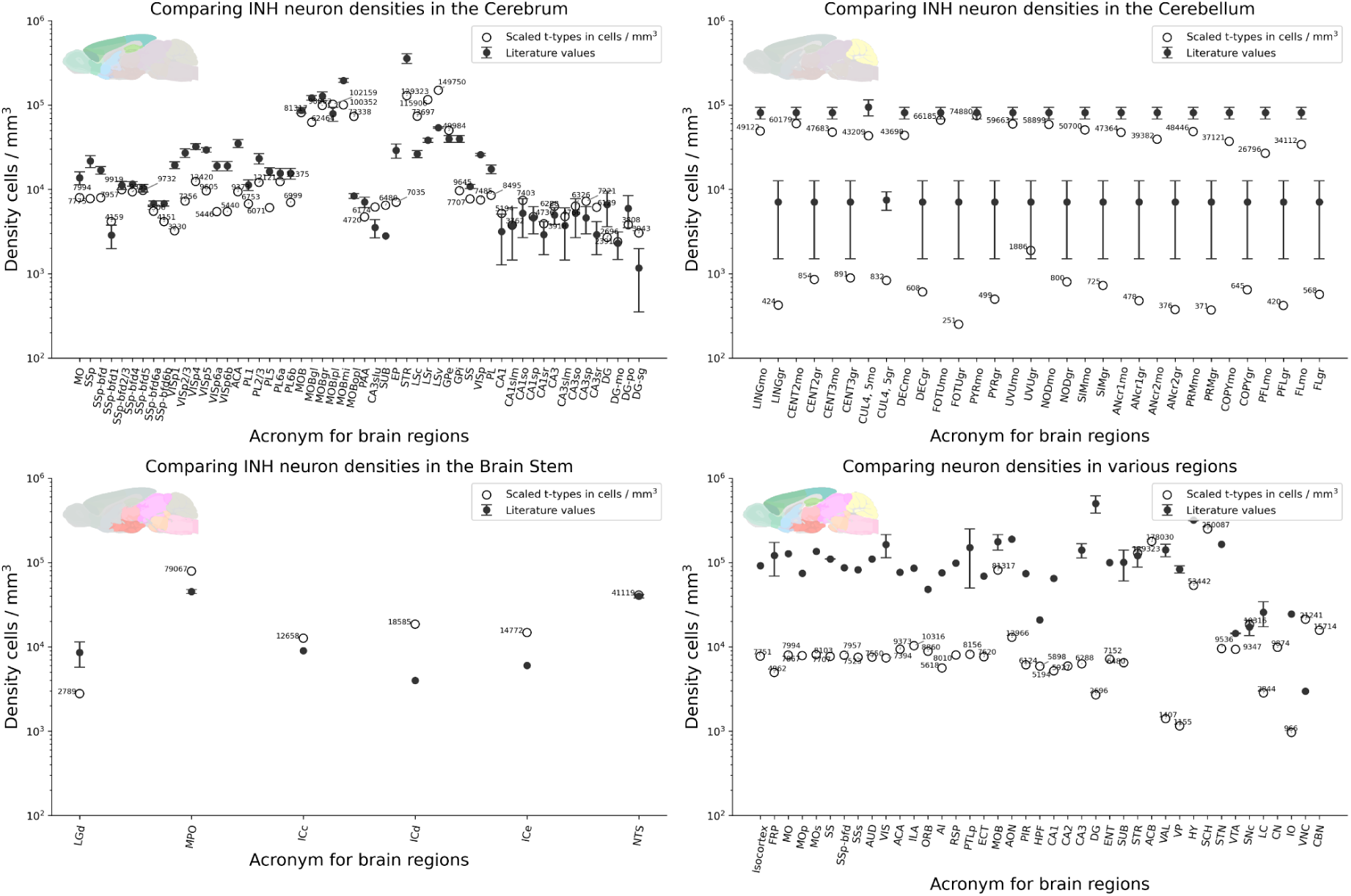
Comparison of predicted scaled density with literature values. **A.** We extracted inhibitory neuron densities from the Cerebrum from the scaled atlas and compared them with inhibitory neuron numbers from the literature. **B.** Extracted from the Cerebellum **C.** Extracted from the brain stem **D.** Comparing total neuron density values from the scaled atlas with values from the literature [7].

## Discussion

This study presents a 3D atlas of estimated transcriptomic, morphological and electrophysiological cell type densities offering a multimodal perspective on diversity in the mouse brain.

We used the complete mouse brain taxonomy and the MERFISH dataset from the ABC Atlas, an extended CCFv3 annotation volume, and an array of corrective and scaling methods to first create a transcriptomic type (t-type) densities atlas. We then leveraged a probabilistic mapping approach to infer morpho-electrical (me-) types from t-types across brain regions.

### Enhancing Interoperability Between Anatomical Annotation Systems

One of the key outcomes of this work is the projection of cell densities into a 3D reference space. The ABC Atlas introduced a hierarchical annotation system for their 3D reference space (CCFv3), where the tiling of parcellations reflects the resolution at which cells were identified, following the point-to-point mapping between the original MERFISH coordinate space and the CCF space. This approach enabled users to aggregate cells based on parcellation assignments at varying anatomical levels. We reconciled differences between this new system (substructures, structures, divisions, categories, organs) and older anatomical annotations (leaf regions, intermediate and higher levels of the hierarchy, called parent regions).

However, none of these anatomical label reference systems are flawless. Some regions are either absent from the set of annotations or exist as annotations without assigned voxels. Additionally, certain voxels could not be mapped to the lowest level of the hierarchy, leading to inconsistencies when calculating volumes. To address these limitations and ensure optimal interoperability between systems, we utilized the Blue Brain Project’s enhanced CCFv3 annotation volume and reference system (Fig. 2S1). By integrating additional metadata for each cell, users can navigate across different annotation systems. However, this approach comes with biological limitations, particularly in homogenizing regions where positional data was only available at higher levels of the hierarchy during the MERFISH registration process.

Overall, as datasets become increasingly large and complex, users from diverse scientific backgrounds need enhanced interoperability and compatibility.

### Addressing Limitations in MERFISH Cell Density Estimations: Scaling Techniques

One key finding of this work is that MERFISH slices struggle to accurately capture cellular densities in dense brain regions. Specifically, in areas such as the granular layers of the cerebellum, the olfactory bulb, and the hippocampus, we observed empty spaces between identified cells, where one would expect higher cell concentrations. This limitation is consistent across all slices. To address this, we introduced a series of scaling techniques to better model total cell numbers across the brain, in some cases increasing the estimated cell counts by up to twenty-fold. Similar to the annotation system, we took a balanced approach by offering multiple scaling methods to adjust the original cell counts. We compared unscaled and scaled cell numbers with available literature values and developed a strategy for transplanting densities that accommodates a wide range of applications, research communities, and computational modeling efforts.

Another important finding is that MERFISH-derived transcriptomic densities yield a different total cell count in the mouse brain compared to the neuron scaling rules for primate brains [5]. While certain cell-dense regions were found to have fewer cells than expected due to technical limitations, the majority of the brain exhibited significantly higher cell counts. These results support more recent results like the Blue Brain Project’s Mouse Cell Atlas [7,47] and from the Allen Brain Atlas [48–50]. These atlases have utilized advanced imaging and molecular labeling techniques to generate more granular data, revealing greater regional variability in cell densities. For instance, the Blue Brain Project’s Cell Atlas [47], with its whole-brain dataset and refined alignment algorithms, suggests higher localized neuron densities, especially in specific cortical layers and subcortical regions. Similarly, the Brain Initiative Cell Census atlases and other Nissl-stained slice datasets show local variations in cell densities that deviate from the scaling rule’s averages, highlighting how methodological advancements are uncovering more complexity in brain composition than previous scaling models suggested. High-resolution scRNA-seq and advanced stereological counting techniques have further revealed more intricate distributions of both neuronal and non-neuronal cell types, demonstrating higher cell densities, particularly in regions with specialized functions [49,51].

#### Re-evaluating Neuronal Populations in White Matter from MERFISH and High-Resolution Imaging

One of the important findings from the generated neuron densities was the significant presence of cell bodies in white matter regions. Upon examining the QC values from [8], we found no significant differences between white and gray matter areas, nor between neuronal and non-neuronal cell types in the white matter. Additionally, we assessed the positions of neuronal cell bodies and found no significant clustering near the boundaries between white and gray matter, suggesting no misalignment in the registration process to the reference atlas.In contrast, in brain regions adjacent to large white matter areas, which are more prone to bleed-over effects, the correct ratio of excitatory and inhibitory neurons, along with non-neuronal cells, was observed.

Several studies have also highlighted the presence of notable neuronal populations in the mouse brain white matter, challenging previous assumptions. For example, research has identified specific densities of NeuN-positive neurons in white matter regions, such as the cingulum and external capsule, with some studies revealing even higher densities in genetically modified models relevant to neurological conditions [52].

Additionally, high-resolution electron microscopy studies have identified features of neuronal activity and axonal density, contributing to the growing understanding of white matter’s role beyond just being a conduit for neural communication. These newer approaches to cell counting, imaging, and genetic analysis indicate that neuron numbers in specific white matter tracts might indeed be higher than previously expected. This has prompted a closer re-evaluation of neuronal counts in mouse brain white matter [53–55].

### Probabilistic Mapping and Cross-modal Validation

Our initial validation steps suggest that the probabilistic mapping preserves broad distinctions among cell types across transcriptomic, electrophysiological, and morphological modalities (Fig. 1S1). The alignment between patch-seq and reference t-type labels demonstrated satisfactory functional consistency, especially for major classifications, such as excitatory versus inhibitory neurons. This consistency aligns with previously reported mappings e.g. [28] and suggests that the mapping approach maintains fundamental cell-type characteristics. However, it should be noted that the validation of morphological alignment relied on qualitative assessments due to limited quantitative data.

### Challenges in Extrapolating Me-types to Uncovered T-types

One of the central goals of this approach was to generalize me-type predictions to t-types not represented in the patch-seq data. By identifying predictive genes for m-types and e-types while minimizing regional variability, we generated a subspace that allowed for extrapolation to other brain regions (Fig.1 C and D). Our optimization of gene counts and neighbor parameters for kNN analysis provided a robust basis for this extrapolation.

However, due to the inherent variability in transcriptomic profiles across brain regions, this generalization may oversimplify certain cellular complexities, limiting predictive accuracy. Although this approach holds potential for extending cellular characterizations, the probabilistic extrapolations require further validation. Additionally, we propose a method to customize the me-type composition for specific regions by incorporating special types defined by the user (see Supplementary Methods: *Special Regions Case*). This approach leverages expert knowledge to mitigate the limitations of the current generalization framework and enhance the biological relevance of the resulting composition.

### Limitations of mRNA Density Estimations from Me-types

Our comparison of mRNA densities derived from t-types and me-types highlights both the utility and limitations of the probabilistic approach. While mRNA densities inferred from me-types were relatively uniform across brain regions, the reduction from 4,804 t-types to 458 me-types introduced some homogeneity that obscures finer transcriptomic distinctions (Fig 4. B and C). This reduction led to a relative error, especially for gene-specific counts, though overall regional compositions were less affected. These findings suggest that the model may adequately capture general expression trends but lacks the specificity to accurately reflect individual gene expression patterns. Therefore, this approach may be better suited for broad overviews of cell-type distributions rather than precise transcriptomic reconstructions.

### Regional Homogeneity in Cell-type Compositions: me-types vs. t-types

The clustering of brain regions by me-type and t-type compositions illustrates the trade-offs of using a reduced framework for cell-type representation. Although t-type compositions showed higher specificity in distinguishing brain regions, clustering based on me-types effectively condensed cell-type profiles, resulting in more generalized brain region groupings (Fig. 5). The identification of six clusters for me-type composition compared to eight for t-types indicates that me-type-based groupings may overlook subtle variations between brain regions. However, the relatively small difference in cluster numbers suggests that, despite the reduction in complexity, the me-type framework is still biologically relevant.

In conclusion, this study presents a probabilistic framework for creating a 3D atlas of t-, m-, and e-type densities, offering insights into cell-type diversity across the mouse brain. Our approach effectively integrates multimodal data and produces biologically relevant regional density estimates. The atlas and pipeline provide a flexible, scalable tool for exploring large-scale cell-type distributions, with potential applications in computational modeling and translational neuroscience.

## Data Records

The source data from the AIBS is available here: https://allen-brain-cell-atlas.s3.us-west-2.amazonaws.com/index.html

As the data is constantly growing the main portal’s website is: https://portal.brain-map.org/atlases-and-data/bkp/abc-atlas

The original publication on the ABC Atlas can be found here:

Yao, Z., van Velthoven, C.T.J., Kunst, M. *et al.* A high-resolution transcriptomic and spatial atlas of cell types in the whole mouse brain. *Nature* 624, 317–332 (2023). <u>10.1038/s41586-023-06812-z</u>

The source data from patch-seq experiments can be found from theri original publications [27,28].

## Code availability

The software used in this study is publicly available: https://github.com/BlueBrain/Molsys-transcriptomic-atlas https://github.com/YannRoussel/probabilistic_mapping_extention

Example code and directions: https://github.com/cveraszto/TME-types

https://github.com/YannRoussel/probabilistic_mapping_extention/blob/main/README.md

## Acknowledgements

We thank Karin Holm for editing assistance and for comments on the manuscript, and Darshan Mandge for e-type information. We are grateful to Sébastien Piluso for providing the extended annotation.

## Supplementary Methods

### File format

To store cell type densities, one can easily access every parcellation for the presence (or absence) of its cell types. It is also trivial to look at each cell type and list all parcellations where they are present. The difference compared to Yao et al. is the estimated density values in cells / mm^3^. For ease of use we generated 3D atlases for every cell type on the cluster level using the extended CCFv3 annotation volume. For any given region the voxels inherit the average cell density of the region. The implicit knowledge of the cell type atlas is that the absence of a cell type is similarly strong information. Thus to make calculations simple, we set voxels outside of the brain to np.nan, marking a clear difference between not present and not possible. The resulting 3D arrays are stored as nrrd files. For in-between calculations, e.g. for cell type / region information which are stored in a dict of dataframes we stored the information in pickle format.

Total cell counts were calculated for every region by multiplying the density values with the region’s total voxel count. This information is useful for comparing them with literature values where total cell counts were estimated. Storing this information in a single pickle file is the most convenient.

Finally, once density information is available in every leaf region, larger regions (higher in the anatomical hierarchy) can be readily calculated by summing up leaf regions which are part of a larger brain structure. This way, we can estimate densities in all levels of the anatomical hierarchy for all cell types, and larger cell groups, e.g. inhibitory, excitatory neurons or non-neuronal cells and store it as a table in a csv file.

### CCFv3 - Annotation volumes

The mouse brain reference atlas or CCFv3 annotation volume, derived from 1,675 mouse brains, provides the ideal template for a 3D mouse brain atlas. Its smooth surface and expert validation make it one of the best and most up-to-date resources in the field. However, to ensure flexibility and future developments, we designed the pipeline in such a way that any type of cell atlas can be used as a template to generate a 3D atlas populated with comprehensive density data. To create a 3D brain representation of every cell type we used the extended version of the CCFv3 annotation volume (Fig. 2S1). This comes with the added benefit of providing coverage of the entire brain. This atlas has more complete versions of both the olfactory bulb, the cerebellum, and the medulla, while it still adheres to the hierarchical structure of the common coordinate framework version 3 from the AIBS. As the rostral dimensions are higher, the 0 origin coordinate is shifted by 350 μm compared to the AIBS CCFv3 annotation volume.

Average cell densities were projected into the reference atlas space, and each voxel inherited a density value for any given cell type, based on the average value of all data from the brain slices covering the region the voxel belonged to. We used the 25 μm^3^ voxel size version to speed up calculations. However, as the pipeline calculates average densities, one can use any version of the annotation volume or any 3D representation of the brain to generate 3D cell atlases, as long as it can match the region ids or the region names, or the parcellations from the AIBS.

Dealing with 5000+ clusters can be daunting. Using the ABC Atlas transcriptomic hierarchy we could combine clusters and their densities into subclasses, supertypes or classes. Since many literature values are available on the classes level, we summed up cluster densities into neuron, inhibitory-, excitatory-, modulatory-, glia (astr+oligo+microglia), astrocyte, oligodendrocyte, microglia densities. This data can be stored in 3D format as nrrd files. Average cell type densities are then easily calculated for the entire annotation hierarchy for validation purposes. For any region in the hierarchy can be broken down to its constituent leaf regions. Cell densities could be summed up and neatly stored in a single dataframe.

### Special regions case

The pipeline allows for the overwriting of cortical me-types with specialized me-types in specific brain regions (e.g., Cerebellum) when the expected densities for these unique me-types are available. The goal of this customization is to identify optimal groupings of t-types, following their hierarchical classification, so that the densities of the grouped t-types best approximate the expected densities of these specialized me-types. This is achieved by defining the me-type densities as:

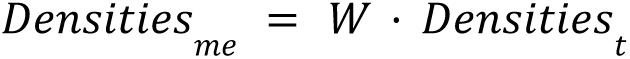

where *W* represents probabilistic weights that are optimized to minimize the distance between the computed me-type densities and the expected densities for specialized regions.

To implement this, hierarchical clustering of t-types is performed based on their expression profiles, creating clusters that preserve similarities among t-types. These clusters are then used to compute aggregated densities, which are normalized and compared to the custom me-type densities via Euclidean distance. The probabilistic weights *W* are iteratively adjusted to minimize this distance, ensuring that the derived me-type densities for the specialized regions align closely with the known, expected densities. This method provides a flexible way to integrate region-specific information into the probabilistic map, improving the accuracy of the density estimates for non-cortical brain regions while maintaining consistency with the overall pipeline.

## Supplementary material

**Supplementary table 1:** Leaf regions not covered by MERFISH slices. 3 out 9 regions are from the grey matter, the Bed nucleus of the anterior commissure is from the Pallidum (Cerebrum), while Frontal pole, layer 1, and Retrosplenial area, dorsal part, layer 4 are from the Isocortex (Cerebrum). The other 6 regions are from the brain’s white matter.

| id | name | closest region |
| --- | --- | --- |
| 68 | Frontal pole, layer 1 | Frontal pole, cerebral cortex |
| 545 | Retrosplenial area, dorsal part, layer 4 | Retrosplenial area, dorsal part |
| 380 | cuneate fascicle | dorsal column |
| 198 | pyramidal decussation | corticospinal tract |
| 1051 | direct tectospinal pathway | tectospinal pathway |
| 287 | Bed nucleus of the anterior commissure | Pallidum, caudal region |
| 1039 | Gracile nucleus | Dorsal column nuclei |
| 949 | vomeronasal nerve | cranial nerves |
| 164 | central canal, spinal cord/medulla | ventricular systems |

### m-type list

**m-type list:**
|  |  |  |
| --- | --- | --- |
| IN_dend_0_ax_0 | IN_dend_3_ax_6 | IN_dend_7_ax_4 |
| IN_dend_0_ax_2 | IN_dend_3_ax_7 | IN_dend_7_ax_7 |
| IN_dend_0_ax_3 | IN_dend_3_ax_8 | IN_dend_7_ax_8 |
| IN_dend_0_ax_4 | IN_dend_3_ax_9 | IN_dend_8_ax_1 |
| IN_dend_0_ax_5 | IN_dend_4_ax_2 | IN_dend_8_ax_7 |
| IN_dend_0_ax_6 | IN_dend_4_ax_3 | IN_dend_8_ax_8 |
| IN_dend_0_ax_8 | IN_dend_4_ax_5 | IN_dend_8_ax_9 |
| IN_dend_0_ax_9 | IN_dend_4_ax_6 | IN_dend_9_ax_0 |
| IN_dend_1_ax_0 | IN_dend_4_ax_8 | IN_dend_9_ax_1 |
| IN_dend_1_ax_1 | IN_dend_4_ax_9 | IN_dend_9_ax_2 |
| IN_dend_1_ax_2 | IN_dend_5_ax_0 | IN_dend_9_ax_3 |
| IN_dend_1_ax_3 | IN_dend_5_ax_3 | IN_dend_9_ax_4 |
| IN_dend_1_ax_4 | IN_dend_5_ax_4 | IN_dend_9_ax_5 |
| IN_dend_1_ax_5 | IN_dend_5_ax_5 | IN_dend_9_ax_8 |
| IN_dend_1_ax_7 | IN_dend_5_ax_6 | IN_dend_9_ax_9 |
| IN_dend_1_ax_8 | IN_dend_5_ax_7 |  |
| IN_dend_1_ax_9 | IN_dend_5_ax_8 | PC_dend_0_ax_8 |
| IN_dend_2_ax_0 | IN_dend_5_ax_9 | PC_dend_1_ax_7 |
| IN_dend_2_ax_2 | IN_dend_6_ax_0 | PC_dend_2_ax_4 |
| IN_dend_2_ax_3 | IN_dend_6_ax_2 | PC_dend_2_ax_7 |
| IN_dend_2_ax_4 | IN_dend_6_ax_3 | PC_dend_2_ax_8 |
| IN_dend_2_ax_5 | IN_dend_6_ax_7 | PC_dend_3_ax_0 |
| IN_dend_2_ax_6 | IN_dend_6_ax_8 | PC_dend_3_ax_4 |
| IN_dend_2_ax_9 | IN_dend_6_ax_9 | PC_dend_3_ax_6 |
| IN_dend_3_ax_0 | IN_dend_7_ax_0 | PC_dend_3_ax_7 |
| IN_dend_3_ax_2 | IN_dend_7_ax_1 | PC_dend_3_ax_8 |
| IN_dend_3_ax_3 | IN_dend_7_ax_2 | PC_dend_8_ax_4 |
| IN_dend_3_ax_5 | IN_dend_7_ax_3 | PC_dend_8_ax_7 |
|  |  | PC_dend_8_ax_8 |

### e-types list

bAC : burst accommodating

bIR : burst irregular

bNAC : burst non-accommodating

bSTUT : burst stuttering

cAC : continuous accommodating

cADpyr : continuous adapting pyramidal neurons

cIR : continuous irregular

cNAC : continuous non-accommodating

cSTUT : continuous stuttering

dNAC : delayed non-accommodating

dSTUT : delayed stuttering

